# A theoretical approach for quantifying the impact of changes in effective population size and expression level on the rate of coding sequence evolution

**DOI:** 10.1101/2021.01.13.426437

**Authors:** T. Latrille, N. Lartillot

**Affiliations:** Université de Lyon, Université Lyon 1, CNRS, Laboratoire de Biométrie et Biologie Évolutive UMR 5558, F-69622 Villeurbanne, France; École Normale Supérieure de Lyon, Université de Lyon, Université Lyon 1, Lyon, France

**Keywords:** protein stability, substitution rate, population-genetics, drift, expression level

## Abstract

Molecular sequences are shaped by selection, where the strength of selection relative to drift is determined by effective population size (*N*_e_). Populations with high *N*_e_ are expected to undergo stronger purifying selection, and consequently to show a lower substitution rate for selected mutations relative to the substitution rate for neutral mutations (*ω*). However, computational models based on biophysics of protein stability have suggested that *ω* can also be independent of *N*_e_, a result proven under general conditions. Together, the response of *ω* to changes in *N*_e_ depends on the specific mapping from sequence to fitness. Importantly, an increase in protein expression level has been found empirically to result in decrease of *ω*, an observation predicted by theoretical models assuming selection for protein stability. Here, we derive a theoretical approximation for the response of *ω* to changes in *N*_e_ and expression level, under an explicit genotype-phenotype-fitness map. The method is generally valid for additive traits and log-concave fitness functions. We applied these results to protein undergoing selection for their conformational stability and corroborate out findings with simulations under more complex models. We predict a weak response of *ω* to changes in either *N*_e_ or expression level, which are interchangeable. Based on empirical data, we propose that fitness based on the conformational stability may not be a sufficient mechanism to explain the empirically observed variation in *ω* across species. Other aspects of protein biophysics might be explored, such as protein-protein interactions, which can lead to a stronger response of *ω* to changes in *N*_e_.

## 1 Introduction

Molecular sequences differ across species due to the particular history of nucleotide substitutions along their respective lineages. These substitutions in turn are the result of the interplay between evolutionary forces such as mutation and selection, whose relative forces are determined by the amount of random genetic drift. These forces have effects at different levels: mutations are carried by molecular sequences, selection is mediated at the level of individuals, while random genetic drift is a population sampling effect. Yet, they jointly contribute to the long-term molecular evolutionary process. Thus, the challenge of the study of molecular evolution is to tease out their respective contributions, based on comparative analyses.

One main aspect of this challenge is to correctly evaluate the role of random drift in the long term evolutionary process. Population genetics theory implies that the strength of drift, due to the stochastic sampling of mutations, is less pronounced in lineages with large effective population size (*N*_e_), and as a consequence, the purification by selection of weakly deleterious mutations is more effective in large populations. This fundamental idea is at the core of the nearly-neutral theory of evolution. This theory posits that a substantial fraction of mutations are deleterious or weakly deleterious, and as a result, predicts that the substitution rate (relative to the neutral expectation), called *ω*, decreases along lineages with higher *N*_e_ (Ohta, 1972, 1992).

This prediction has been more quantitatively examined under the assumption that the selective effects of mutations are drawn from a fixed distribution of fitness effects (DFE) (Kimura, 1979; Welch *et al.*, 2008). Assuming a gamma distribution for the DFE, a key result obtained in this context is an approximate allometric scaling of *ω* as a function of *N*_e_ (i.e. 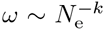), where *k* is the shape parameter of the DFE. In practice, DFEs are strongly leptokurtic, which thus predicts a weak negative relation between *ω* and *N*_e_.

The study of protein-coding sequences evolution fostered another modelling approach, based on genotype-fitness maps instead of distribution of fitness effects. In this alternative approach, the selective effect of a mutation depends on the fitness of both the source and the target amino acids involved in the mutation event (Halpern and Bruno, 1998; Rodrigue *et al.*, 2010; Tamuri and Goldstein, 2012). Even though this modelling approach differs substantially from the one assuming a fixed DFE, it also predicts a negative correlation between *ω* and *N*_e_, at least when the process is at equilibrium (Spielman and Wilke, 2015; Dos Reis, 2015).

Conversely, one striking theoretical result was the proof that *ω* is in fact predicted to be independent of *N*_e_ under relatively general circumstances, namely, whenever (i) the fitness is a log-concave function of a phenotype and (ii) the phenotype itself is equimutable. Equimutability states that the distribution of phenotypic changes due to mutation is independent of the current phenotype of individuals (Cherry, 1998). This general theoretical argument has been invoked in the context of *in silico* experiments of protein sequence evolution, assuming that proteins are under selection for their thermodynamic stability, with fitness being proportional to the folding probability of the protein (Goldstein, 2013). Thermodynamic stability is itself computed using a 3D structural model of the protein. These computational experiments have led to the observation that *ω* is essentially independent of *N*_e_. An explanation proposed for this result is that the distribution of changes in free energy of folding (ΔΔ*G*) due to mutations is approximately independent of the current free energy (Δ*G*), thus making the free energy of folding essentially equimutable.

However, the equimutability assumption is a relatively strong one, which also conflicts with combinatorial considerations about the relation between sequence and phenotype (Serohijos *et al.*, 2012). For example, if a protein sequence is already maximally stable, only destabilizing (or neutral) mutations can occur. More generally, assuming that the stability of a protein sequence reflects an underlying fraction of positions having already accepted destabilizing amino acids, then the probability of destabilizing mutational events is in turn expected to directly depend on the current stability of the protein.

Altogether, depending on the theoretical model mapping sequence to fitness, *ω* can be either independent or negatively correlated to *N*_e_, or even positively if considering adaptive evolution and environmental changes (Lanfear *et al.*, 2014).

Empirically, variation in *ω* between lineages has been inferred using phylogenetic codon models applied to empirical sequences (Yang and Nielsen, 1998; Zhang and Nielsen, 2005). Confronting branch-specific *ω* estimates to life-history traits such as body mass or generation time uncovered a positive correlation (Popadin *et al.*, 2007; Nikolaev *et al.*, 2007). Subsequently, integrative inference methods combining molecular sequences and life-history traits have also found that *ω* correlates positively with traits such as longevity and body mass (Lartillot and Poujol, 2011; Figuet *et al.*, 2017). Since lineages with a large body size and extended longevity typically correspond to species with low *N*_e_ (Romiguier *et al.*, 2014), these empirical correlations suggest a negative correlation between *ω* and *N*_e_, thus confirming the theoretical prediction of the nearly-neutral theory of evolution. However, the universality and robustness of the correlation between *ω* and life-history traits is still debated. Results have not been entirely consistent across independent studies. The correlation was found to be either not statistically significant (Lartillot and Delsuc, 2012), or even in the opposite direction depending on the specific clade under study or the potential biases taken into account (Lanfear *et al.*, 2010; Nabholz *et al.*, 2013; Lanfear *et al.*, 2014; Figuet *et al.*, 2016).

If empirical evidence for a negative correlation of *ω* with *N*_e_ is still not totally convincing, another empirical correlation is known to be much more robust. Indeed, expression level or protein abundance is one of the best predictors of *ω*, with highly expressed proteins typically having lower *ω* values, a correlation clearly significant although relatively weak (Duret and Mouchiroud, 2000; Rocha and Danchin, 2004; Drummond *et al.*, 2005; Zhang and Yang, 2015; Song *et al.*, 2017). Theoretical models, also based on protein stability, have been invoked to explain this negative correlation between *ω* and expression level (Wilke and Drummond, 2006; Drummond and Wilke, 2008). According to this argument, selection against protein misfolding due to toxicity, which is stronger for more abundant proteins, induces abundant proteins to evolve toward greater stability, resulting in a more constrained and more slowly evolving protein coding sequence (Serohijos *et al.*, 2012).

The possibility that expression level and *N*_e_ might play similar roles in the evolution of proteins has already been noticed. More precisely, under models of selection against protein misfolding, the free energy of folding Δ*G* is predicted to vary similarly along a gradient of either *N*_e_ or expression level (Serohijos *et al.*, 2013). As a corollary, under strict equimutability of Δ*G*, these computational models imply that *ω* should also be independent of expression level (Serohijos *et al.*, 2012), akin to what is predicted with regards to changes in *N*_e_.

Altogether, both theoretical results and empirical analyses are not yet conclusive about the question of how *ω* depends on *N*_e_ and expression level. In particular, the theoretical response of *ω* to changes in both *N*_e_ and expression level has not been quantified and, most importantly, has not been related to the specific map between genotype, phenotype and fitness. Such an analytical development would be useful to more decisively confront the theoretical predictions relating *ω* to both *N*_e_ and expression level to empirical data. Ultimately, relating proteins structural parameters to the response of *ω* would help to bridge the gap between protein thermodynamics on one side and comparative genomics on the other side.

Lastly, the theoretical results discussed so far are valid only at the mutation-selection-drift balance. In a non-equilibrium regime, however, and at least under a model assuming a site-independent genotype-fitness map, an increase in *N*_e_ first leads to an increase in *ω* caused by adaptive substitutions, and subsequently a decrease in *ω* due to stronger purifying selection in the long term (Jones *et al.*, 2016). Studying only equilibrium properties can thus be misleading. For this reason, the dynamic response of *ω* to changes in *N*_e_ must also be addressed, quantified, and its connection with the underlying selective landscape better characterized. Dynamic properties of *ω* to changes in *N*_e_ are of theoretical interest but are also empirically relevant, such that, if overlooked they could thwart the relation between theoretical expectations and empirical estimates.

In this context, the aim of the present study is to characterize the dynamics and equilibrium response of *ω* to changes in *N*_e_ and expression level, and to relate this response to structural parameters of the model. To this effect, we develop a general mathematical approach to derive a quantitative approximation of the response of *ω* to changes in *N*_e_ and expression level, in the context of a given genotype-phenotype-fitness map, as depicted in figure 1. In the light of previously published empirical estimates from protein thermodynamics and comparative genomics, we discuss the articulation between empirical data and our mechanistic model. We also discuss some of the alternative biophysical mechanisms that could determine the selective landscape on protein-coding sequences, and how they would modulate the response of *ω* to changes in *N*_e_ and expression level.

**Figure 1:**
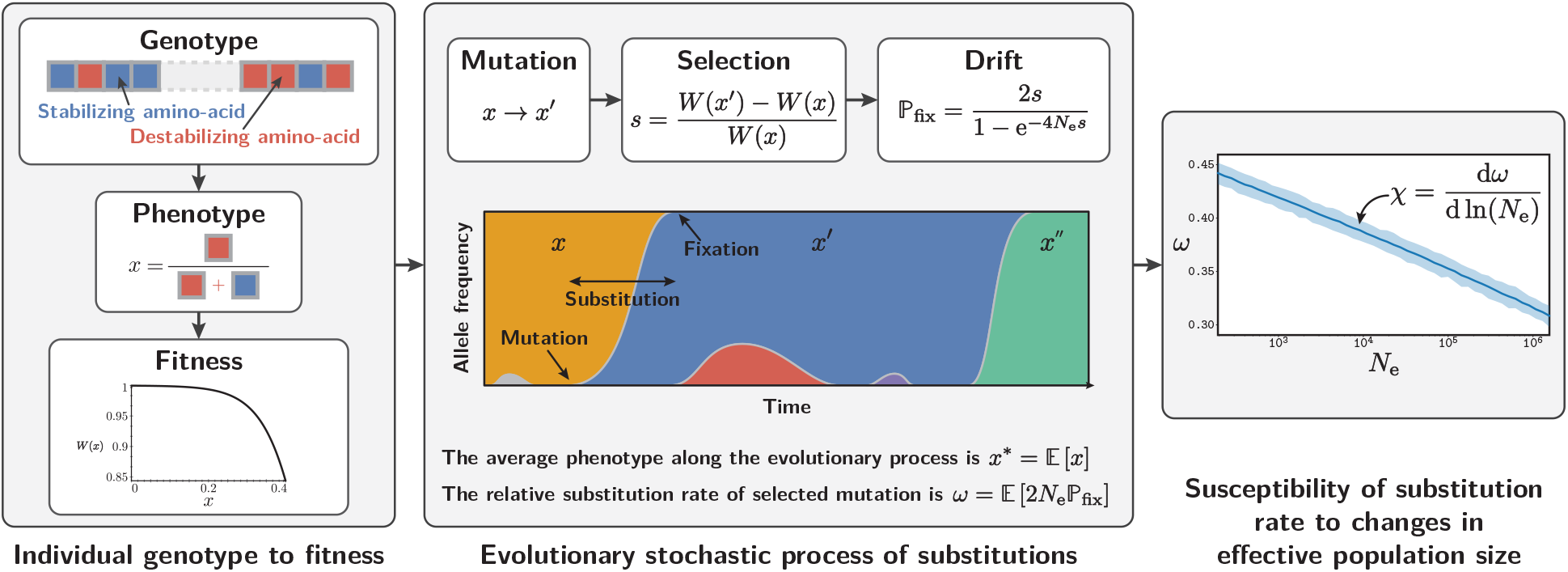
Outline of the theoretical results. The genotype to fitness relationship is depicted in the left panel. The phenotype (*x*) is a real-valued function of the genotype (i.e. the amino-acid sequence), and is defined in our model as the fraction of destabilizing amino acids in the sequence. Fitness is a decreasing log-concave function of the phenotype, depending on structural parameters of the model. Once the relation from genotype to fitness is defined, the substitution process proceeds as presented in the middle panel. For a given effective population size *N*_e_, the evolutionary process results in an average value of the phenotype *x*\* and an average substitution rate (relative to the neutral rate) *ω*. Averaging over time is equivalent to determining the statistical equilibrium, by ergodicity of the stochastic process. The slope of the scaling of the equilibrium *ω* as a function of log-*N*_e_ defines the susceptibility *χ*, which is a function of the structural parameters defined by the phenotype-fitness map.

## 2 Results

### 2.1 Models of evolution

The results that are presented below are valid for a general category of models of sequence evolution, based on an additive trait *x*, such that the coding positions of the sequence contribute additively to the trait. The trait is under directional selection specified by a decreasing and log-concave fitness function *W* (*x*). As a specific example, we more specifically consider a model of protein evolution under the constraint of thermodynamic stability, as depicted in the left panel of figure 1. This model is inspired from previous work (Williams *et al.*, 2006; Goldstein, 2011; Pollock *et al.*, 2012), except that we make several simplifying assumptions, allowing us to derive analytical equations.

In the original biophysical model, protein stability is determined by the difference in free energy between the folded and unfolded conformations, called Δ*G* and measured in kcal/mol. Technically, free energy is computed based on the 3D conformation of the protein and using statistical potentials. As a result, the stabilizing or destabilizing effect of an amino acid at a particular site depends on amino acids present in the vicinity in 3D conformation, thus implementing what has been called specific epistasis (Starr and Thornton, 2016).

Here, we approximate this model such that the (de-)stabilizing effect at a particular site, such as measured by the ΔΔ*G* of the mutation, does not depend on other neighbouring residues, thus disregarding specific epistasis (Dasmeh *et al.*, 2014). Instead, each site contributes independently and additively to Δ*G*. In addition, we assume that, for each site of the sequence, only one amino acid is stabilizing the protein. All 19 other amino acids are equally destabilizing. Each site bearing a destabilizing amino acid contributes an excess of ΔΔ*G* > 0 (in kcal/mol) to the total Δ*G*. The smallest achievable value of Δ*G*, obtained when all amino acids of the sequence are stabilizing, is noted Δ*G*_min_ < 0. In this model, the most succinct phenotype of a given genotype (i.e. sequence) is just the proportion of destabilizing amino acids in the sequence, defined as 0 ≤ *x* ≤ 1. Thus Δ*G* is a linear function of *x*:

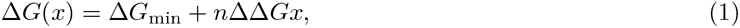

where *n* is the number of sites in the sequence.

For a given Δ*G*, thermodynamic equations allows one to derive the proportion of protein molecules that are in the native (folded) conformation in the cytoplasm. This fraction is assumed to be a proxy for fitness, motivated in part by the fact that a protein must be folded to perform its function. A slightly different model will be considered below, in order to take into account protein expression level (see section 2.3).

Analytically, the fitness function is given by the Fermi Dirac distribution and is typically close to 1, leading to a first-order approximation (Goldstein, 2011):

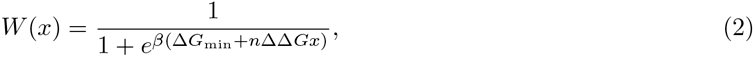

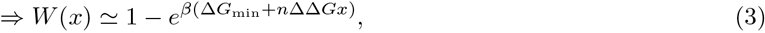

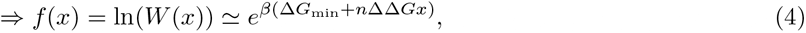

where *W* is the Wrightian fitness for a given phenotype and *f* is the Malthusian fitness (or log-fitness). Here, Δ*G*_min_ and ΔΔ*G* are defined as above, and the parameter *β* is 1.686 mol/kcal at 25°*C* (or 298.2K).

Of note, even though the phenotypic effect of a mutation at a given site does not depend on the amino-acids that are present at other sites (i.e. the trait is additive), the fitness effect of a mutation still depends on other sites (i.e. the log-fitness is not additive). As a result, the molecular evolutionary process is site-interdependent, a property referred to as non-specific epistasis (Starr and Thornton, 2016; Dasmeh and Serohijos, 2018).

### 2.2 Response of *ω* to changes in *N*_e_. Analytical approximation

For a given effective population size *N*_e_, the evolutionary process reaches an equilibrium (figure 1, middle panel). This substitution rate at this equilibrium, normalized by the substitution rate of neutral of mutations to discard the influence of the underlying mutation rate, is denoted *ω*. This relative rate can also be interpreted as the mean fixation probability of mutations scaled by the fixation probability of neutral alleles *p* = 1/2*N*_e_, the mean being weighted by the probability of occurrence of mutations in the population. As a result, an *ω* < 1 indicates that mutations are negatively selected on average, and *ω* decreases with increasing strength of purifying selection.

In this section we present an analytical approximate solution for the response of *ω* after a change in *N*_e_ (in log space), as depicted in the right panel of figure 1. We call this response the susceptibility of *ω* to changes in *N*_e_, and denote it as *χ*:

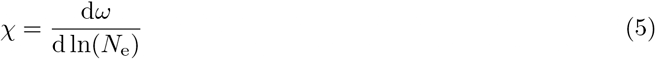

Deriving *χ* is done in two steps. First, we determine the mean phenotype at equilibrium, when evolutionary forces of mutation, selection and genetic drift compensate each other. Subsequently, differential calculus is used to compute the response of the equilibrium phenotype to a change in *N*_e_, which allows us to ultimately derive an equation for *χ*. The main results of our derivation are given both in the general case of any (log-concave) phenotype-fitness map, and in the specific case of the biophysical model introduced above. A more detailed derivation is available in the supplementary materials.

For a given genotype, mutations can have various effects: they can increase or decrease the proportion of destabilizing amino acids, or do nothing if the mutation is between two destabilizing amino acids. To derive the probabilities of such events to occur, we also make the simplifying assumption that all transitions between amino acids are equiprobable. Altogether, any mutation in the sequence can then have a phenotypic effect of 0 or *δx* = 1/*n*, with probabilities of transitions equal to:

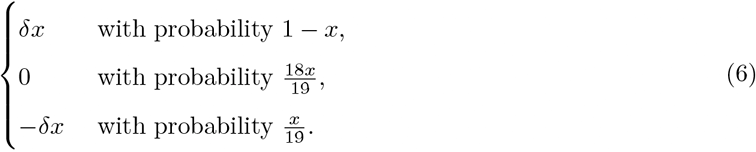

In the extreme case of an optimal phenotype (*x* = 0), only destabilizing mutations are proposed. Moreover, the probability to propose a stabilizing mutation (effect −*δx*), or a neutral mutation (effect 0), is proportional to *x*. Conversely, the probability to propose a destabilizing mutation is equal to (1 − *x*). As a result, the mutation bias is proportional to (1 − *x*)/*x*. This mutation bias fundamentally reflects a combinatorial effect, due to the number of mutational opportunities available in either direction.

Second, we need to determine the strength of selection acting on mutations. Destabilizing mutations are selected against with a negative selection coefficient which can be approximated by:

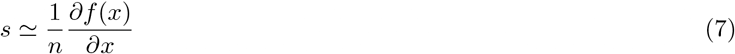

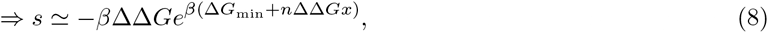

where *f* = ln(*W*) is the log-fitness (or Malthusian fitness). Conversely, stabilizing mutations will be under positive selection with opposite sign but same absolute value. It is important to realize that the selective effect is dependent on *x*. Furthermore, because the fitness function is log-concave, the absolute value of *s* increases with *x*.

Based on these expressions for the mutational and selective pressures, one can then study the trajectory followed by the evolutionary process. Starting from an optimal sequence, mostly destabilizing mutations will occur, some of which may reach fixation and accumulate until selection coefficients against new deleterious mutations is too strong, at which point the protein will reach a point of equilibrium called marginal stability (Taverna and Goldstein, 2002; Bloom *et al.*, 2007). Most importantly, the probability of fixation of mutations is affected by genetic drift, and thus depends on the effective population size (*N*_e_). At the equilibrium between mutation, selection and drift, the process fluctuates through the occurrence of advantageous and deleterious substitutions compensating each other. This equilibrium can be determined by expressing the constraint that the selection coefficient of substitutions is expected to be null on average (Goldstein, 2013). Formally, and after simplification, the equilibrium phenotype denoted *x*\* is given in the general case by:

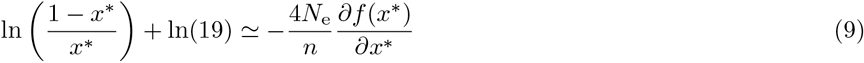

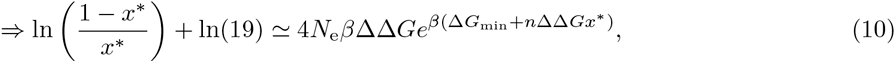

in the more specific case of the biophysical model. This equation essentially expresses the mutation-selection equilibrium: the left-hand side of the equation is the log of the mutation bias at *x*, while the right-hand side is simply 4*N*_e_*s*, the scaled selection coefficient.

This equation cannot be solved explicitly for *x*\*, but a qualitative intuition on the consequences of change in *N*_e_ to the equilibrium phenotype *x*\* is given in figure 2. Intuitively, an increase in *N*_e_ results in a more optimal phenotype, closer to 0. The mutation bias (left-hand side of equation 10) decreases with *x* while the strength of selection (right-hand side of equation 10) increases with *x*, and the equilibrium phenotype is obtained at their intersection. An increase in *N*_e_ leads to shifting the selective response upward, which then results in a leftward shift of the equilibrium phenotype (i.e. closer to 0). The leftward shift is smaller for selective strengths characterized by a steeper curve, resulting in qualitatively weaker susceptibility of the equilibrium phenotype to changes in *N*_e_

**Figure 2:**
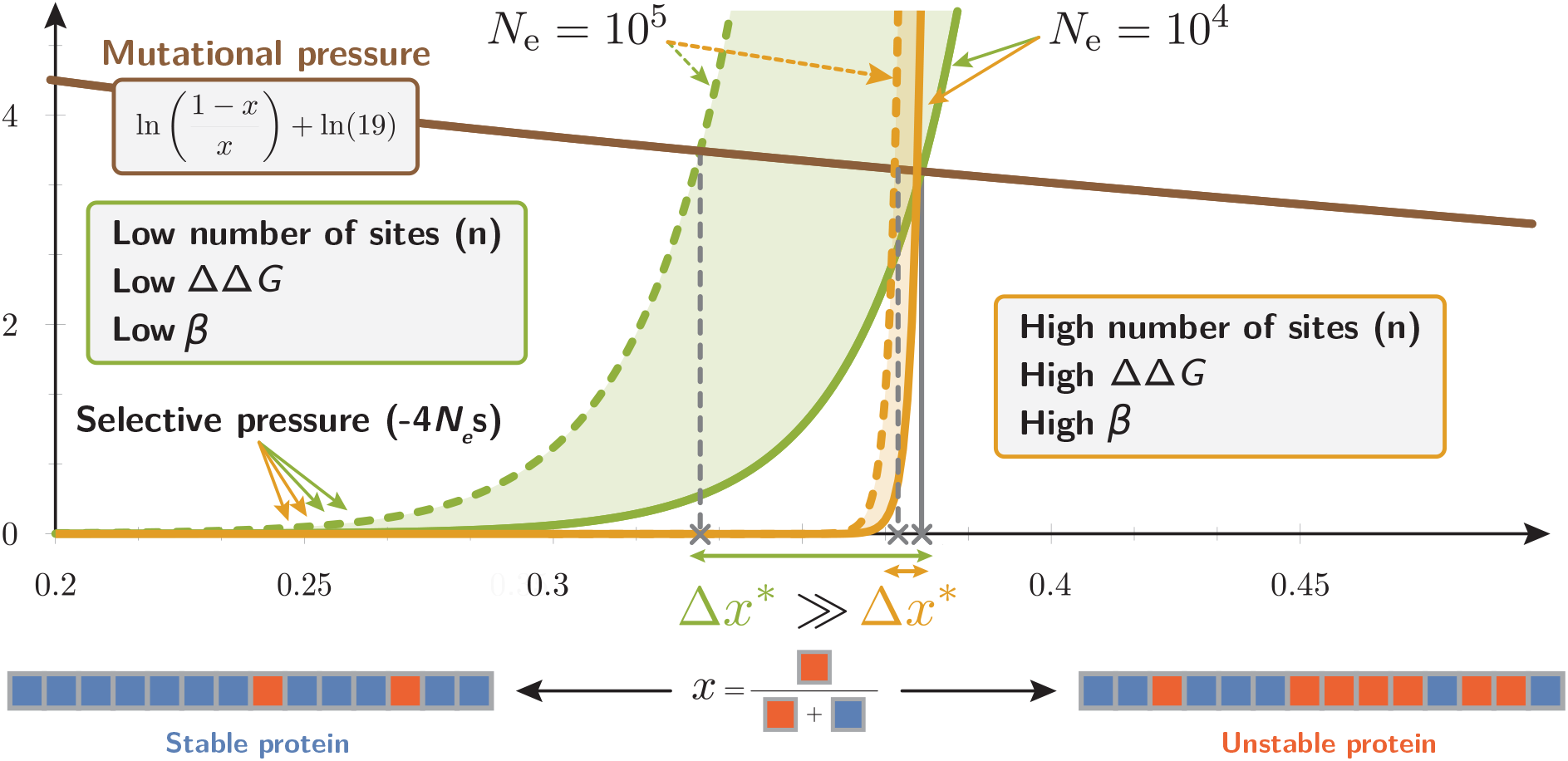
Response of the equilibrium phenotype after a change in *N*_e_. The equilibrium phenotype *x*\* is obtained when the selective pressure equals to the mutational pressure (equation 10). The selective pressure (right-hand side of eq. 10) increases exponentially with *x* where *βn*ΔΔ*G* is the exponential growth rate (yellow and green curves). When *βn*ΔΔ*G* is large, increasing *N*_e_ by an order of magnitude (yellow dotted curves) very moderately impacts the equilibrium phenotype (small Δ*x*\*). In contrast, for small *βn*ΔΔ*G* (green curves), the equilibrium phenotype is more strongly impacted by a change in *N*_e_ (large Δ*x*\*). Finally, response of *x*\* to changes in *N*_e_ reflects the response of *ω* since both are approximately equal (equation 11).

The results obtained thus far only relate the equilibrium phenotype (*x*\*) to *N*_e_. To capture how *ω* varies with *N*_e_, we also need to obtain an expression for *ω* as a function of *x*\*. At equilibrium we can derive (supplementary materials) the expected substitution rate of mutations, and thus *ω*, which simply approximates to:

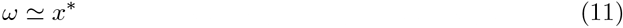

This simple approximation is due to the fact that the substitutions between two destabilizing amino acids (which are neutral) compose the largest proportion of proposed mutations having a substantial probability of fixation (equation 6). In contrast, stabilizing mutations are rare, while destabilizing mutations have a low probability of fixation. Since there is a fraction *x*\* of sites already occupied by a destabilizing amino-acid, these neutral substitutions occur at rate *x*\*.

Combined together, these analytical approximations yield the susceptibility (equation 5) of *ω* to a change in *N*_e_:

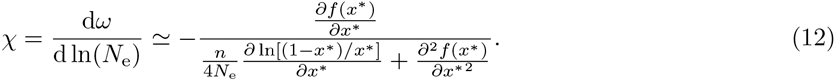

The two terms of the denominator correspond to the derivative of the mutational bias and the scaled selection coefficient, respectively. However, the mutational bias decreases weakly with *x* (blue curve on figure 2) while the strength of selection increases sharply with *x* (red and green curves). As a consequence, the derivative of the mutational bias is much lower than the derivative of the selection coefficient around the equilibrium point (i.e. the phenotype is *nearly* equimutable). The second term can therefore be ignored, which leads to a very compact equation for susceptibility *χ* in the general case:

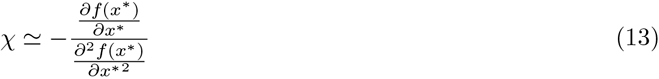

The susceptibility is thus equal to the inverse of the relative curvature, i.e. the ratio of the second to the first derivatives, of the log-fitness function, taken at the equilibrium phenotype. Of note, this susceptibility is strictly negative for decreasing log-concave fitness functions, asserting that *ω* is a decreasing function of *N*_e_. In addition, the susceptibility itself is low in absolute value (i.e. *ω* responds more weakly) for strongly concave log-fitness functions. This equation quantitatively captures the intuition developed in figure 2, namely that the response of *ω* is very weak if the selection curve is very steep around the equilibrium set point (red curve compared to green curve).

In the specific case of the biophysical model, the susceptibility (*χ*) further simplifies to:

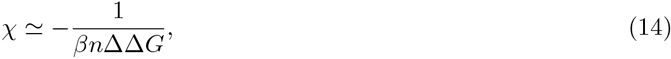

meaning that *ω* is linearly decreasing with *N*_e_ (in log scale) since *χ* is independent of *x*\*, or, in other words, that the exact value of the equilibrium phenotype has no impact on the slope. Moreover, only the compound parameter *β*ΔΔ*Gn* has an impact on the slope of the linear relationship. Thus, in particular, the slope of the linear relationship between *ω* and *N*_e_ is affected by ΔΔ*G* but not by Δ*G*_min_. Of note, empirically, only relative values of *N*_e_ (up to a multiplicative constant) are required to obtain an estimate of *χ*.

### 2.3 Response of *ω* to changes in protein expression level

Effective population size is not the sole predictor of *ω*, and expression level (or protein abundance) is also negatively correlated to *ω*. However, our previous model, which assumes that fitness is proportional to the folded fraction, and is thus independent of protein abundance, does not express the fact that selection is typically stronger for proteins characterized by higher levels of expression. An alternative biophysical model is to assume that each misfolded protein molecule has the same relative effect on fitness, caused by its toxicity for the cell (Drummond *et al.*, 2005; Wilke and Drummond, 2006; Drummond and Wilke, 2008; Serohijos *et al.*, 2012).

Our general derivation can be directly applied to this case. For a given protein with expression level *y* and a cost *A* representing the selective cost per misfolded molecule (positive constant), the fitness and selection coefficient can be defined as follows:

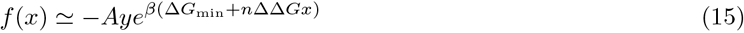

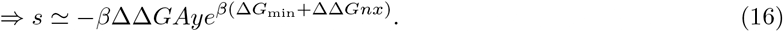

Under this model, the total selective cost of a destabilizing mutation is now directly proportional to the total amount of misfolded proteins. This fitness function leads to the following expression for the mutation-selection-drift equilibrium:

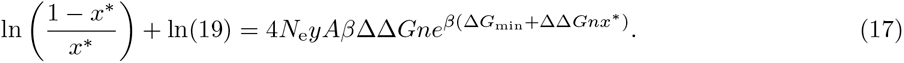

Importantly, in this equation, *N*_e_ and *y* are confounded factors appearing only as a product. This means that increasing either *N*_e_ or *y* leads to same change in equilibrium phenotype, and hence the same change in *ω*. In other words, the susceptibility of the response to changes in either *N*_e_ or expression level is the same:

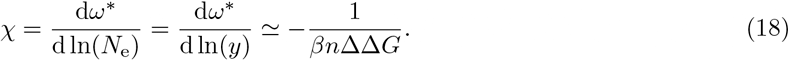

A similar result can be obtained under other models relating phenotype to fitness, for example if the selective cost is due to translational errors (supplementary materials). Alternatively if the protein is assumed to be regulated such as to reach a specific level of functional protein abundance under a general cost-benefit argument (Cherry, 2010; Gout *et al.*, 2010), a multiplicative factor depending solely on the expression level is prefixed (supplementary materials). Altogether, we theoretically obtain the same linear decrease of *ω* with regards to either effective population size or expression level (in log space) under a broad variety of hypotheses.

### 2.4 Simulation experiments

Our theoretical derivation of the susceptibility of *ω* to changes in *N*_e_ (and expression level) is based on several simplifying assumptions about the evolutionary model and makes multiple approximations. In order to test the robustness of our main result, we therefore conducted systematic simulation experiments, relaxing several of these assumptions. In each case, simulations were conducted under a broad range of values of *N*_e_, monitoring the average *ω* observed at equilibrium and plotting the scaling of these measured equilibrium *ω* as a function of *N*_e_.

Specifically, with respect to mutations, our derivation assumes that all amino-acid transitions are equiprobable, or in other words, the complexity of the genetic code is not taken into account. Simulating evolution of DNA sequence and invoking a matrix of mutation rates between nucleotide allows us to test the robustness of our results to this assumption. Furthermore, with regard to the phenotypic effects of amino-acid changes, in our derivation, we assumed that all destabilizing amino acids have an identical impact on protein stability. In reality, one would expect conservative amino-acid replacements to be less destabilizing than radical changes.

This assumption is relaxed in our simulation, such that destabilizing mutations in each position are now proportional to the Grantham distance (Grantham, 1974) between the optimal amino acid in this position and the amino acid proposed by the non-synonymous mutation. Finally, our derivation assumes that the number of sites in the sequence (*n*) is large, such that the selection coefficient is well approximated by the fitness derivative (equation 7). The robustness of this approximation was tested by conducting simulations with finite sequences of realistic length (*n* = 300 coding positions).

These simulation experiments demonstrate, first, that the relation between *ω* and log-*N*_e_ is indeed linear, at least in the range explored here, and that the slope of the linear regression matches the expected theoretical value (figure 3A). Secondly, we observe that the parameter Δ*G*_min_ has virtually no effect on the slope of the linear regression, as also expected theoretically (figure 3B). Instead, decreasing Δ*G*_min_ (to more negative values) merely results in an overall increase in *ω* over the whole range of *N*_e_ (i.e. has an impact on the intercept, not on the slope of the relation). This is due to the fact that decreasing Δ*G*_min_ shifts the equilibrium to higher *x*\*, since more destabilizing sites can then reach fixation before reaching the point of marginal stability.

**Figure 3:**
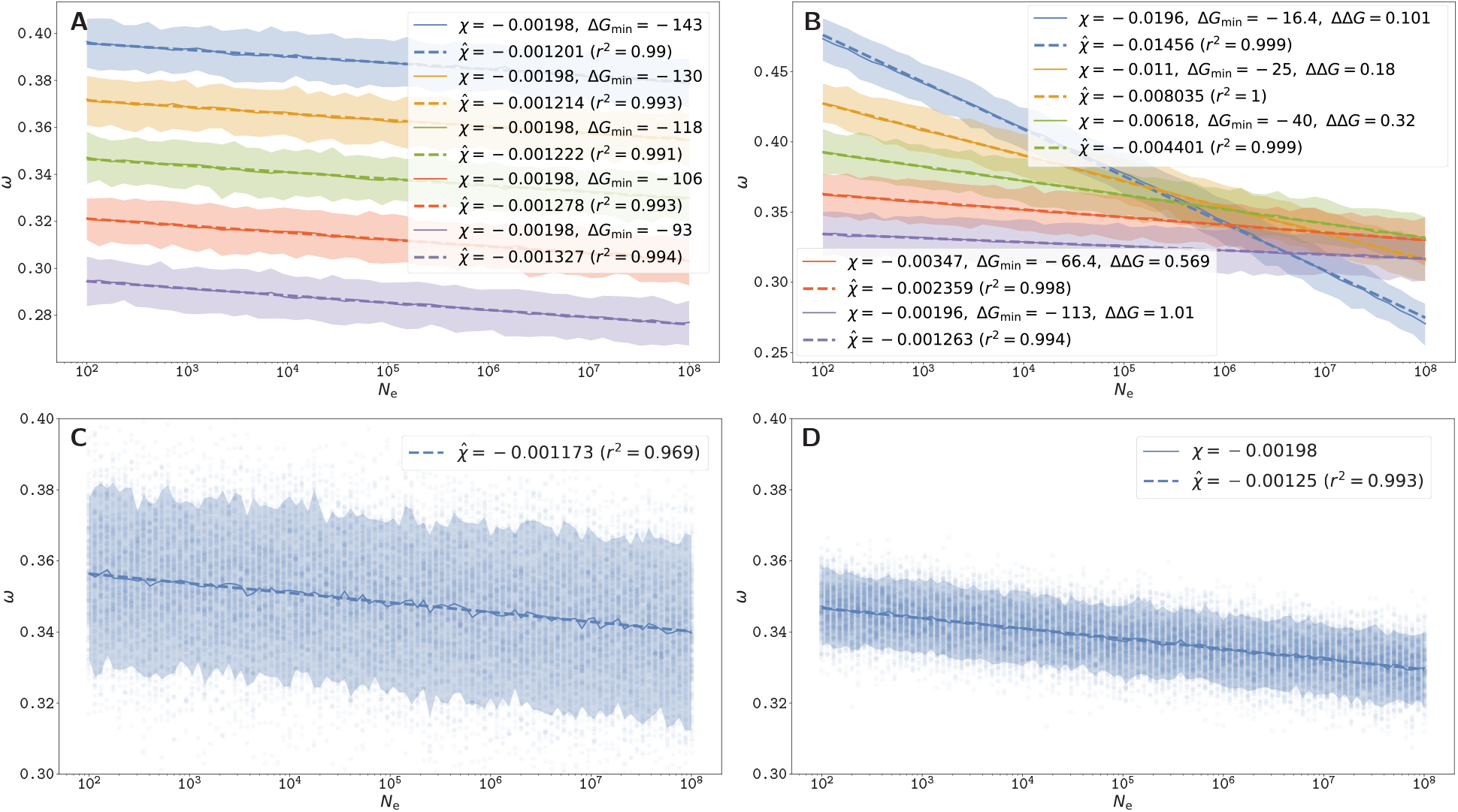
Scaling of equilibrium *ω* as a function of log-*N e*, under the additive phenotype model using Grantham distances (A,B,D) or the explicit biophysical model using a statistical potential (C), with *n* = 300 and *β* = 1.686. 200 replicates per *N*_e_ value are shown (dots). Solid lines are average over replicates, and shaded areas are 90% confidence interval. The slope (or susceptibility 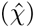, is estimated by linear regression (dashed lines). (A): Δ*G*_min_ are given in the legend, and ΔΔ*G* = 1. Decreasing Δ*G*_min_ (to more negative values) increases *ω* but does not impact the slope. (B): ΔΔ*G* is increased and Δ*G*_min_ is changed accordingly such that the equilibrium value *x*\* is kept constant, by solving numerically equation 10. The estimated susceptibility 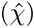 decreases proportionally to the inverse of ΔΔ*G*, as predicted by our theoretical model. (C): Stability of the folded native state is computed using 3D structural conformations and pairwise contact potentials. (D): Additive model with Δ*G*_min_ = *−*118 kcal/mol and ΔΔ*G* = 1 kcal/mol matches structural model shown in C (although with less variance).

Finally, we relaxed our assumption that each site of the sequence contributes independently to Δ*G*, by taking into account the 3D structure of protein and using a statistical potential to estimate Δ*G* (supplementary materials). We implemented the original model considered in Williams *et al.* (2006), Goldstein (2011) and Pollock *et al.* (2012), in which the free energy is computed based on the 3D conformation using pairwise contact potential energies between neighbouring amino-acid residues (Miyazawa and Jernigan, 1985). The original works showed that under this model, *ω* is approximately independent of *N*_e_ (Goldstein, 2013). Using extensive simulations in order to obtain sufficient resolution, we observe that *ω* is in fact weakly dependent on *N*_e_, being again approximately linear with log-*N*_e_ (figure 3C). Moreover, the observed slope 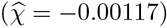 matches the slope obtained under the model of additive Δ*G* (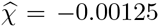, figure 3D), considering an empirical ΔΔ*G* = 1.0 kcal/mol for destabilizing mutations and *n* = 300. In this experiment (figure 3D), Δ*G*_min_ was set to *−*118 kcal/mol, which is the Δ*G* of the optimal (maximally stable) sequence of 300 sites (Goldstein, 2011).

### 2.5 Time to relaxation

Although the equilibrium value of *ω* after changes in *N*_e_ is an important feature of the *ω*-*N*_e_ relationship, another characteristic that is scarcely studied is the dynamic aspect (Jones *et al.*, 2016), particularly the relaxation time to reach the new equilibrium *ω*. We observed in our simulations that the determining factor of the relaxation time is the number of sites *n* (figure 4A), such that the return to equilibrium is faster for longer sequences. This observation matches the theoretical prediction that more mutational opportunities are available for longer sequences, driving the trait close to equilibrium at a faster rate.

**Figure 4:**
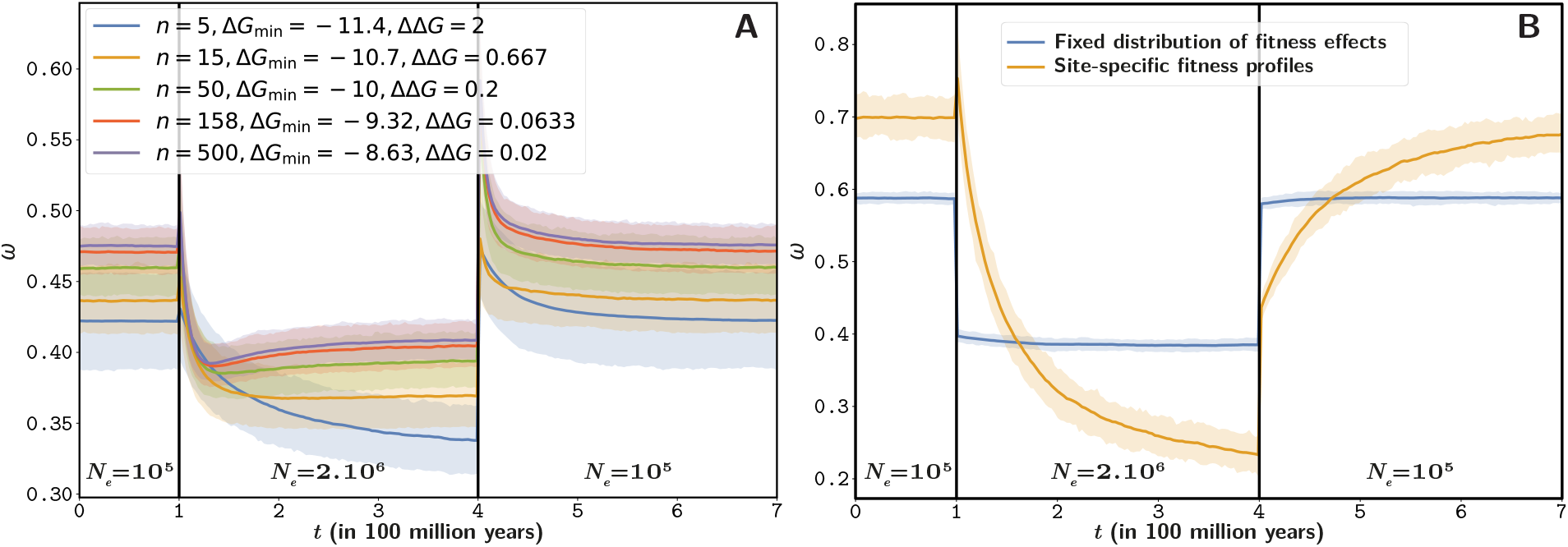
Relaxation of *ω* after a change in *N*_e_ Solid line corresponds to the average over 1000 replicates and the shaded area corresponds to the 90% interval among replicates. The mutation rate (*μ*) is 1*e−*8 per year per site, and the total evolutionary period is 700 million years. (A): *β* = 1.686 for all simulations. The DNA sequence of 500 sites is divided into exons of equal size. However the number of sites per exon changes between simulations from *n* = 5 to *n* = 500. Moreover, ΔΔ*G* is changed according to the exon size such that *n*ΔΔ*G* (and as a result, the susceptibility) are kept constant, and Δ*G*_min_ is changed accordingly such that the equilibrium value *x*\* is kept constant, by solving numerically equation 10. Thus, regardless of exon size, *x*\* and *χ* are kept constant and thus the observed effect is due to the number of sites in the exon. We observe that increasing the number of sites leads to a reduced time to reach the new equilibrium. (B): In the context of a time-independent fitness landscape (yellow curve), where each amino acid has different fitness (site-specific profiles), the time taken to reach the new equilibrium value of *ω* after a change in *N*_e_ is long. In the context of a fixed distribution of fitness effects (blue curve), the relaxation time is non-existent and the new equilibrium value of *ω* is reached instantaneously.

It may be useful to compare the relaxation pattern observed here with the predictions under two alternative models of sequence evolution, representing two extreme scenarios. On one hand, having fitness modelled at the level of sites, such as contemplated by many phylogenetic mutation-selection models (Halpern and Bruno, 1998; Rodrigue *et al.*, 2010; Tamuri and Goldstein, 2012), leads to a situation where every site has to adapt on its own to the new change in *N*_e_. The relaxation time is then very long, on the order of the inverse of the per-site substitution rate. On the other hand, assuming a fixed distribution of fitness effect (DFE) as in Welch *et al.* (2008), the response of *ω* is instantaneous (figure 4B). Our model is effectively in between these two extreme scenarios.

Another characteristic observed in these non-equilibrium experiments is the discontinuity of *ω* after a change in *N*_e_. Most importantly, both an increase and decrease in *N*_e_ lead to a discontinuity (figure 4A & 4B). These non-equilibrium behaviors can both be explained mechanistically. Under low *N*_e_, the phenotype is far away from the optimal phenotype because the efficacy of selection is weaker. A sudden increase in *N*_e_ results first in a short traction toward a more optimal phenotype, which results in a suddenly higher *ω*, caused by a transient adaptation of the protein toward a higher stability. Conversely, under high *N*_e_ the phenotype is closer to optimal and the purification of deleterious mutations is stronger. The reaction to a decrease in *N*_e_ is a relaxation of the purification and thus an *ω* closer to the neutral case, which results into higher *ω* until reaching the point of marginal stability. To note, an increase in *N*_e_ can theoretically and possibly lead to an *ω* that is temporarily greater than 1 due to adaptive evolution (Jones *et al.*, 2016), while a decrease in *N*_e_ always imply an *ω* < 1, as it gives at most a neutral regime of relaxed selection.

## 3 Discussion

We provide a compact analytical result for the equilibrium response (which, by analogy with thermodynamics, we call the susceptibility) of *ω* to changes in *N*_e_, and we relate this response to the parameterization of the genotype-phenotype-fitness map. An application to a model of selection against protein misfolding shows that the response of *ω* to variation in *N*_e_ (in log space) is linear, with a negative slope. Furthermore, this application demonstrates that effective population size and protein expression level are interchangeable with respect to their impact on the response of *ω*. Our compact theoretical results, which were obtained by making several simplifying assumptions, are supported by more complex simulations of protein evolution relaxing these assumptions. In particular, our theoretical predictions are verified under a numerical model of protein evolution in which the free energy is computed based on the 3D structure.

Overall, the susceptibility (*χ*) is a function of the structural parameters of the protein and takes a very simple analytical form, being inversely proportional to the product of three terms: the sequence size, the inverse temperature (*β*), and the average change in conformational energy of destabilizing mutations (ΔΔ*G*). Quantitatively, this product can be several orders of magnitude greater than 1 in practice, such that the susceptibility of *ω*, which is its inverse, is typically small. Previous studies using this model presented an apparent lack of response of *ω* to changes in *N*_e_ (Goldstein, 2013). We refine this result, by observing that there is in fact a very subtle and weak relation, which requires extensive computation to be detected, but which is well predicted by our theoretical derivation. Based on empirical estimates of the structural parameters *β* = 1.686, *n* = 300 sites and ΔΔ*G* = 1.0 kcal/mol for destabilizing mutations (Zeldovich *et al.*, 2007), the estimated susceptibility is 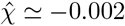. In other words, for a relative increase in *N*_e_ or expression level of 6 orders of magnitude, a factor approximately equal to 0.01 is subtracted from *ω*, a subtle relationship that requires laborious effort to be detected in simulated data.

### 3.1 Adequacy to empirical data

Empirically, variation in *ω* along the branches of phylogenetic trees has been inferred and correlated to proxies of *N*_e_, such as body size or other life-history traits. These analyses showed mitigated support for a negative relation between *ω* and *N*_e_ (Lanfear *et al.*, 2014). More recently, phylogenetic integrative methods refined the estimate of covariation between *ω* and *N*_e_ along lineages by leveraging polymorphism data (Brevet and Lartillot, 2019). This approach gives an estimate of 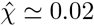 in primates (supplementary materials) at least one order of magnitude greater than the quantitative estimate obtained above from the biophysical model. More empirical data across different clades would be required to robustly consolidate such empirical estimates, but as of yet, these results are challenging the idea of a very weak response.

The relation between *ω* and expression level provides an independent, and potentially more robust, source of empirical observation. Our theoretical results suggest that, under relatively general conditions, the response of *ω* to expression level should be of the same magnitude than the response to *N*_e_. Empirically, the protein expression level is one of the best predictors of *ω* and the empirical estimation of *χ* in fungi, archaea and bacteria varies in the range [−0.046; −0.021] (supplementary materials) extracted from Zhang and Yang (2015). Estimation in animals and plants gives somewhat lower estimates, in the range of [−0.026; −0.004], although still higher (in absolute value) than −0.002.

Additionally, another empirical observation is the negative relation between the mean destabilizing effect of mutations (mean ΔΔ*G*) and the Δ*G* of the protein. Such a relation is empirically observed in Serohijos *et al.* (2012), where the slope of the linear regression is −0.13 (*r*^2^ = 0.04). The slope of the linear correlation observed in our simulations is weaker, with an observed slope of −0.01 (*r*^2^ = 0.29) under the 3D biophysical model, and −0.003 (*r*^2^ = 0.33) under the model of additive phenotype parameterized by ΔΔ*G* = 1 and *n* = 300 (supplementary materials). This observation also sheds light on the correlation between *ω* and *N*_e_ in empirical data and in our model. Indeed, equimutability, or namely that the distribution of ΔΔ*G* of mutations is independent of Δ*G* is a necessary condition to observe independence between *ω* and *N*_e_ (Cherry, 1998). In our model, the average ΔΔ*G* of mutations at equilibrium depends on Δ*G* due to combinatorial considerations, but this dependence is weaker than empirically observed, which also translates into a weaker susceptibility of *ω* to changes in *N*_e_ or expression level than empirically observed.

Thus, overall, the response of *ω* to either *N*_e_ or expression level predicted by the biophysical model considered above seems lower than what is empirically observed. There are several possible explanations for this discrepancy. First, the biophysical model might be valid, but the numerical estimates used for *n* or ΔΔ*G* could be inadequate. A ΔΔ*G* of 1.0 kcal/mol for destabilizing mutations seems to correspond to empirical estimates (Zeldovich *et al.*, 2007). On the other hand, the effective number of positions implicated in the trait might be smaller than the total number of residues in the protein. In our model, all positions in the protein can in principle compensate for the destabilizing effect of a mutation at a particular position. In practice, the number of sites susceptible to compensate each other is probably smaller, resulting in a stronger departure from equimutability.

Alternatively, the biophysical model considered here might be too restrictive. Recent empirical studies have provided evidence against the hypothesis that the rate of sequence evolution is driven solely by the toxicity effect of unfolded proteins (Plata and Vitkup, 2017; Razban, 2019; Biesiadecka *et al.*, 2020). Notably, the response of *ω* to changes in expression level has also been found theoretically to arise as a consequence of protein-protein interactions, where protein may either be in free form or engaged in non-specific interactions (Yang *et al.*, 2012; Zhang *et al.*, 2013). In non-specific interactions at the protein surface, stabilizing amino acids are hydrophilic and destabilizing amino acids are hydrophobic, sticking to hydrophobic residues at the surface of other proteins (Dixit and Maslov, 2013; Manhart and Morozov, 2015).

Our theoretical results can be applied more broadly to protein-protein interactions using a mean-field argument (supplementary materials). Fitting this model with empirical structural estimates (Janin, 1995; Zhang *et al.*, 2008), we obtain a susceptibility of *χ* ≃ −0.2 thus a much stronger response than under the model based on conformational stability. This much stronger response is due to fewer sites in the protein being involved in protein-protein interaction than for conformational stability, in addition to a lower free energy engaged in contact between residues.

Altogether, fitness based on protein stability is a compelling model of molecular evolution, but may not be a sufficiently comprehensive model to explain the amplitude of variation of *ω* empirically observed along a gradient of either effective population size or protein expression level. The net response of *ω* to changes in *N*_e_ or expression level could have several biophysical causes, which in the end would imply a weak but still empirically measurable response.

### 3.2 The statistical mechanics of molecular evolution

This study describes the signature imprinted on DNA sequences by an evolutionary process by merging equations from population genetics and from structural physicochemical first principles. More generally, it outlines a general approach for deriving quantitative predictions about the observable macroscopic properties of the molecular evolutionary process based on an underlying microscopic model of the detailed relation between sequence, phenotype and fitness. In this respect, it borrows from statistical mechanics, attempting approximations to derive analytically tractable results (Sella and Hirsh, 2005; Mustonen and Lässig, 2009; Bastolla *et al.*, 2012, 2017) The robustness of results can be assessed by computational implementations and simulations. Computational models offer a means to test the validity and robustness, while mathematical models offer an intuitive mechanistic mental analogy.

Ultimately, the approach could be generalized to other aspects of the evolutionary process. Beyond *ω*, other macroscopic observables could be of interest, for example site entropy, i.e. the effective number of observed amino acids per site at equilibrium (Goldstein and Pollock, 2016; Jimenez *et al.*, 2018; Jiang *et al.*, 2018), or the nucleotide or amino-acid composition. In addition to *N*_e_, other evolutionary forces could also be considered, for instance the mutational bias or GC-biased gene conversion. The susceptibility of the macroscopic observables to changes in the strength of these underlying forces could then more generally be investigated. As such, the framework outlined here could foster a better understanding of observable signatures of the long-term evolutionary process emerging from ecological parameters and molecular physico-chemical first principles, by carefully teasing out the combined effects of mutation, selection and drift.

## 4 Materials & Methods

Protein sequence evolution is simulated under an origin-fixation model (McCandlish and Stoltzfus, 2014), i.e. the whole population is considered monomorphic and only the succession of fixation events is modeled. Given the currently fixed sequence 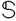, we define 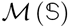 as the set of all possible mutant that are one nucleotide away from 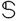. Non-sense mutants are not considered. For a protein of *n* amino-acid sites, 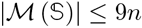, since each codon has a maximum of 9 possible nearest neighbors that are not stop codons. For each mutant sequence 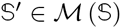, we compute its fitness and subsequently the selection coefficient of the mutant:

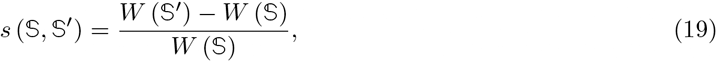

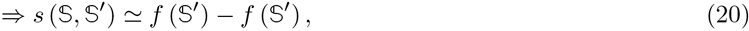

where *W* is the Wrightian fitness for a given phenotype and *f* is the Malthusian fitness (or log-fitness).

The waiting time before the next mutant invading the population, and the specific mutation involved in this event, are chosen using Gillespie’s algorithm (Gillespie, 1977), according to the rates of substitution between 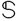 and each 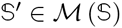, which are given by:

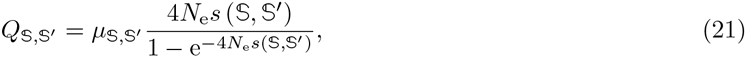

where 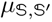 is the mutation rate between 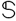 and 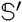, determined by the underlying 4*x*4 nucleotide mutation rate matrix, and 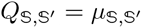 in the case of synonymous substitutions. Various optimizations are implemented to reduce the computation time of mutant fitness. The simulation starts with a burn-in period to reach mutation-selection-drift equilibrium.

### 4.1 Models of the fitness function

Under the additive model for the free energy, the difference in free energy between folded and unfolded state is assumed to be given by:

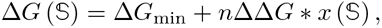

where 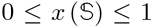 is the distance of 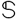 to the optimal sequence (i.e. the fraction of sites occupied by a destabilizing amino-acid). For each site of the sequence, the optimal amino acids are chosen randomly at initialization, and the distance between the current amino acid and the optimal is scaled by the Grantham amino-acid distance (Grantham, 1974). The Wrightian fitness is defined as the probability of our protein to be in the folded state, given by the Fermi-Dirac distribution:

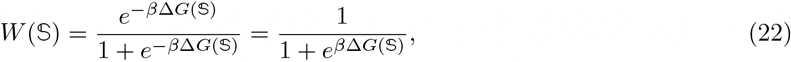

where *β* is the inverse of the temperature (*β* = 1/*kT*).

For simulations under a 3D model of protein conformations, we adapted the model developed in Goldstein and Pollock (2017) to our C++ simulator (see supplementary materials).

For simulations under a site-independent fitness landscape, with site-specific fitness profiles, the protein log-fitness is computed as the sum of amino-acid log-fitness coefficients along the sequence. In this model, each codon site *i* has its own fitness profile, denoted 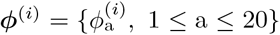, a vector of 20 amino-acid scaled (Wrightian) fitness coefficients. Since 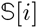 is the codon at site *i*, the encoded amino acid is 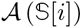, hence the fitness at site *i* is 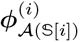. Altogether, the selection coefficient of the mutant 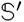 is:

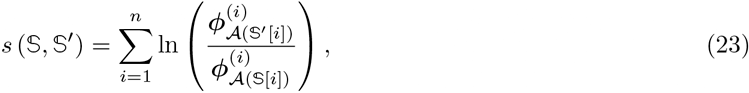

The fitness vectors *φ*^(*i*)^ used in this study are extracted from Bloom (2017). They were experimentally determined by deep mutational scanning.

For simulations assuming a fixed distribution of fitness effects (DFE), the selection coefficient of the mutant 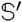 is gamma distributed (shape *k* > 0):

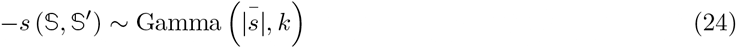

### 4.2 Computing *ω* along the simulation

From the set of mutants 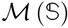 that are one nucleotide away from 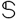, we define the subsets 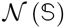 of non-synonymous and synonymous mutants 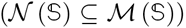. The ratio of non-synonymous over synonymous substitution rates, given the sequence 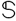 at time *t* is defined as (Spielman and Wilke, 2015; Dos Reis, 2015; Jones *et al.*, 2016):

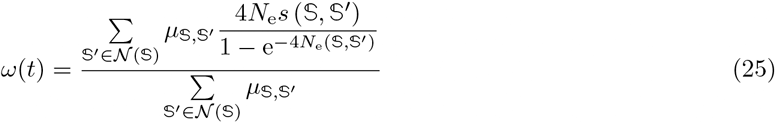

Averaged over all branches of the tree, the average *ω* is:

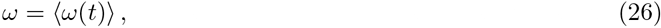

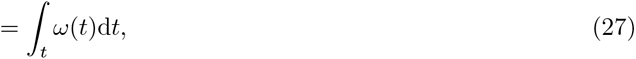

where the integral is taken over all branches of the tree, while the integrand *ω*(*t*) is a piece-wise function changing after every point substitution event.

## 5 Reproducibility - Supplementary Materials

The simulators written in C++ are publicly available under MIT license at https://github.com/ ThibaultLatrille/SimuEvol. The mathematical developments under the general case of an arbitrary additive trait and an arbitrary log-concave fitness function, and the derived susceptibility under various fitness functions, as well as supplementary figures, are available in supplementary materials. The scripts and instructions necessary to reproduce this study are available at https://github.com/ThibaultLatrille/ GenotypePhenotypeFitness.

## 6 Author contributions

TL gathered and formatted the data, developed the new models in SimuEvol and conducted all analyses, in the context of a PhD work (Ecole Normale Superieure de Lyon). TL and NL both contributed to the writing of the manuscript.

## 7 Acknowledgements

We wish to thank Julien Joseph for whiteboard mathematical sessions. We gratefully acknowledge the help of Nicolas Rodrigue and Laurent Duret for their input on this work and their comments on the manuscript. This work was performed using the computing facilities of the CC LBBE/PRABI. Funding: French National Research Agency, Grant ANR-15-CE12-0010-01 / DASIRE.

## Notes

### Competing Interest Statement

The authors have declared no competing interest.

