## Supplementary materials for "A theoretical approach for quantifying the impact of changes in effective population size and expression level on the rate of coding sequence evolution"

**4 Empirical estimation** **10**

**5 Simulation using the 3D structure of protein** **11**

**6 Simulated $\omega$ response to changes in $N_e$** **13**

**7 Simulated relaxation time of $\omega$** **18**

**8 Distribution of fitness effects** **20**

Notations in the main manuscript and the supplementary file are identical, expect that $\Delta\Delta G$ is simplified to $\gamma$ and $\Delta G_{\text{min}}$ is simplified to $\alpha$ in the supplementary (hereby) for formula readability and developments. $\Delta\Delta G$ and $\Delta G_{\text{min}}$ are kept here to refer to the empirical estimations.

### 1 $\omega$ response after a change in $N_e$

#### 1.1 Genotype to phenotype map

Define $n$ as the number of sites in the genotype sequence. Each site can be in one of $\text{K} \geq 2$ states, where only 1 state is defined the stable state, and $\text{K} - 1$ states are unstable. For a given genotype sequence, define phenotype $0 \leq x \leq 1$ as the current proportion of sites in the unstable state. After a mutation, given that only one site can change at a time, the absolute change of $x$ is either 0 or $\delta x = 1/n$. Define $\rho_x(\delta x)$ as the probability to get a change of phenotype equal to $\delta x$, if the current phenotype is $x$:

$$\begin{cases} \delta x & \text{with probability } \rho_x(\delta x) = 1 - x, \\ 0 & \text{with probability } \rho_x(0) = x\left[1 - \frac{1}{\text{K}-1}\right], \\ -\delta x & \text{with probability } \rho_x(-\delta x) = \frac{x}{\text{K}-1}. \end{cases} \tag{1}$$

#### 1.2 Selection coefficient

$s(x, \delta x)$ is the selection coefficient of an effect $\delta x$ if the current phenotype is $x$:

$$
\begin{aligned}
s(x, \delta x) &= \frac{W(x+\delta x) - W(x)}{W(x)}, && (2)\\
&\simeq \frac{1}{W(x)} \frac{\partial W(x)}{\partial x} \delta x, && (3)\\
&\simeq \frac{\partial \ln(W(x))}{\partial x} \delta x, && (4)\\
&\simeq \frac{\partial f(x)}{\partial x} \delta x, && (5)
\end{aligned}
$$

where $W(x)$ is the Wrightian fitness of phenotype $x$, and $f = \ln(W)$ is the log-fitness (or Malthusian fitness). And the selective effect of the opposite change $(-\delta x)$ is the opposite selection coefficient:

$$
\begin{aligned}
s(x, -\delta x) &\simeq -s(x, \delta x) \text{ from eq. 5}, && (6)\\
\iff S(x, -\delta x) &\simeq -S(x, \delta x), && (7)
\end{aligned}
$$

where $S(x^*, \delta x) = 4N_\mathrm{e} s(x^*, \delta x)$ is the scaled selection coefficient.

#### 1.3 Probability of fixation

The probability of fixation of a mutation with effect $\delta x$, for a resident phenotype $x$ is :

$$
\begin{aligned}
\mathbb{P}_\text{fix}(x, \delta x) &= \frac{1 - \mathrm{e}^{-2s(x,\delta x)}}{1 - \mathrm{e}^{-4N_\mathrm{e} s(x,\delta x)}}, && (8)\\
&\simeq \frac{2s(x,\delta x)}{1 - \mathrm{e}^{-4N_\mathrm{e} s(x,\delta x)}}, && (9)\\
&= \frac{2s(x,\delta x)}{1 - \mathrm{e}^{-S(x,\delta x)}}. && (10)
\end{aligned}
$$

And in the case of neutral mutations, the probability of fixation is:

$$
\mathbb{P}_\text{fix}(x, 0) = \frac{1}{2N_\mathrm{e}}. \quad (11)
$$

And the ratio of probability of fixation between selected and neutral mutations is:

$$
\begin{aligned}
\frac{\mathbb{P}_\text{fix}(x, \delta x)}{\mathbb{P}_\text{fix}(x, 0)} &= \frac{2N_\mathrm{e} 2s(x,\delta x)}{1 - \mathrm{e}^{-S(x,\delta x)}} \text{ from eq. 10 and 11}, && (12)\\
&= \frac{S(x,\delta x)}{1 - \mathrm{e}^{-S(x,\delta x)}}. && (13)
\end{aligned}
$$

#### 1.4 Equilibrium phenotype

At equilibrium phenotype $x^*$, the expected selection coefficient of mutation that reached fixation must be 0:

$$
\begin{aligned}
0 &= \mathbb{E}_{\delta x}\left[ s(x^*, \delta x) \mathbb{P}_\text{fix}(x^*, \delta x) \right], && (14)\\
\iff 0 &= \frac{2s(x^*,\delta x)^2}{1 - \mathrm{e}^{-S(x^*,\delta x)}} \rho_{x^*}(\delta x) + s(x^*, 0) \frac{\rho_{x^*}(0)}{2N_\mathrm{e}} + \frac{2s(x^*,-\delta x)^2}{1 - \mathrm{e}^{-S(x^*,-\delta x)}} \rho_{x^*}(-\delta x) \text{ from eq. 10 and 11}, && (15)\\
\implies \frac{2s(x^*,\delta x)^2}{1 - \mathrm{e}^{-S(x^*,\delta x)}} \rho_{x^*}(\delta x) &\simeq \frac{-2s(x^*,\delta x)^2}{1 - \mathrm{e}^{S(x^*,\delta x)}} \rho_{x^*}(-\delta x) \text{ from eq. 7}, && (16)\\
\iff \frac{\rho_{x^*}(\delta x)}{\rho_{x^*}(-\delta x)} &\simeq \mathrm{e}^{-S(x^*,\delta x)} \frac{\mathrm{e}^{-S(x^*,\delta x)} - 1}{\mathrm{e}^{-S(x^*,\delta x)} \left(1 - \mathrm{e}^{S(x^*,\delta x)}\right)}, && (17)\\
\iff \ln\left(\frac{1 - x^*}{x^*}\right) + \ln(\mathrm{K} - 1) &\simeq -S(x^*, \delta x) \text{ from eq. 1}, && (18)\\
\iff \lambda_K(x^*) &\simeq -S(x^*, \delta x), && (19)
\end{aligned}
$$

where $\lambda_K(x^*) = \ln\left(\frac{1-x^*}{x^*}\right) + \ln(\mathrm{K} - 1)$.

##### 1.5 Relative substitution rate ($\omega$) at equilibrium

The substitution rate of all selected relative to the substitution rate of neutral mutations is denoted $\omega$, which can also be interpreted as the mean fixation probability of mutations scaled by the fixation probability of neutral mutations $p = 1/2N_{\mathrm{e}}$.

$$\omega = \mathbb{E}_{\delta x}\left[\frac{\mathbb{P}_{\text{fix}}(x,\delta x)}{\mathbb{P}_{\text{fix}}(x,0)}\right], \tag{20}$$

$$= (1-x)\frac{S(x,\delta x)}{1-\mathrm{e}^{-S(x,\delta x)}} + x\left(\frac{\mathrm{K}-2}{\mathrm{K}-1}\right) + \frac{x}{\mathrm{K}-1}\frac{S(x,-\delta x)}{1-\mathrm{e}^{-S(x,-\delta x)}} \text{ from eq. 1, 10 and 11,} \tag{21}$$

$$= (1-x)\frac{S(x,\delta x)}{1-\mathrm{e}^{-S(x,\delta x)}} - \frac{x}{\mathrm{K}-1}\frac{S(x,\delta x)}{1-\mathrm{e}^{S(x,\delta x)}} + x\left(\frac{\mathrm{K}-2}{\mathrm{K}-1}\right) \text{ from eq. 7.} \tag{22}$$

$\omega^*$ at equilibrium is then determined by the phenotype at equilibrium $x^*$:

$$\omega^* = (1-x^*)\frac{S(x^*,\delta x)}{1-\mathrm{e}^{-S(x^*,\delta x)}} - \frac{x^*}{\mathrm{K}-1}\frac{S(x^*,\delta x)}{1-\mathrm{e}^{S(x^*,\delta x)}} + x^*\left(\frac{\mathrm{K}-2}{\mathrm{K}-1}\right), \tag{23}$$

$$= x^*\left[\frac{2(x^*-1)\lambda_K(x^*)}{\mathrm{K}(x^*-1)+1} + \frac{\mathrm{K}-2}{\mathrm{K}-1}\right] \text{ from eq. 18.} \tag{24}$$

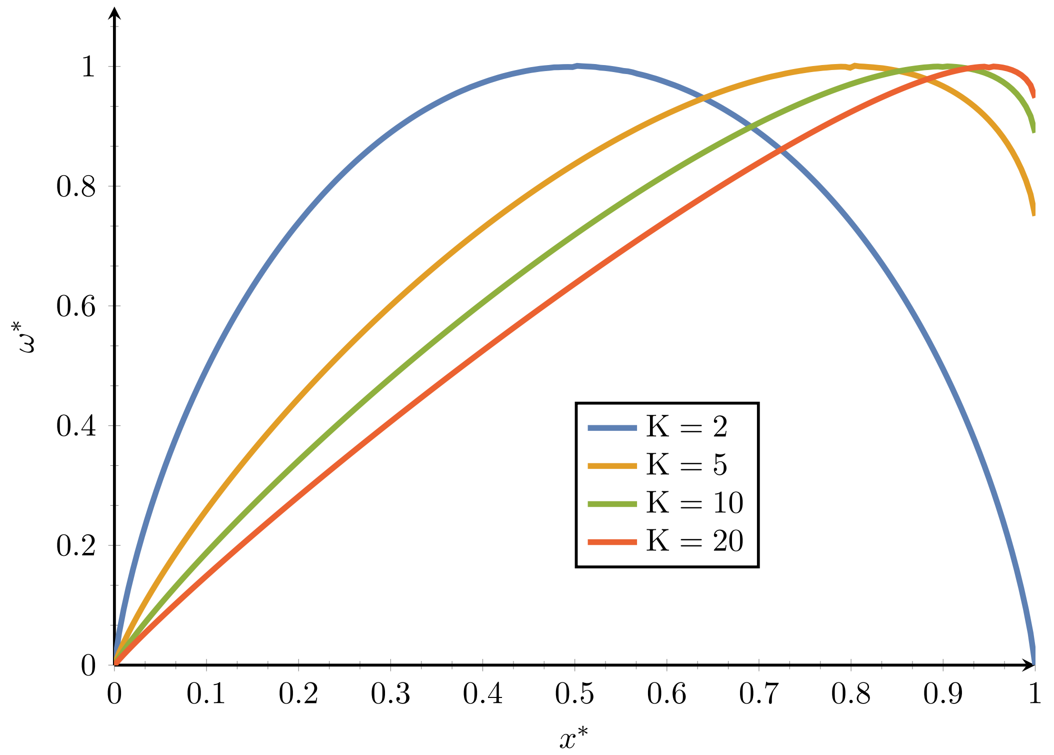

Moreover, given that the number of state if large enough $\mathrm{K} \gg 1$, the equilibrium $\omega$ can be approximated as:

$$\omega^* = x^*\left[\frac{2(x^*-1)\lambda_K(x^*)}{\mathrm{K}(x^*-1)+1} + \frac{\mathrm{K}-2}{\mathrm{K}-1}\right], \tag{25}$$

$$\simeq x^* \tag{26}$$

And the derivative of $\omega^*$ w.r.t to $x^*$ is:

$$\frac{\mathrm{d}\omega^*}{\mathrm{d}x^*} = 2\left[\frac{\mathrm{K}(x^*-1)+1+\left[\mathrm{K}(x^*-1)^2+2x^*-1\right]\lambda_K(x^*)}{(\mathrm{K}(x^*-1)+1)^2}\right] + \frac{\mathrm{K}-2}{\mathrm{K}-1}. \tag{27}$$

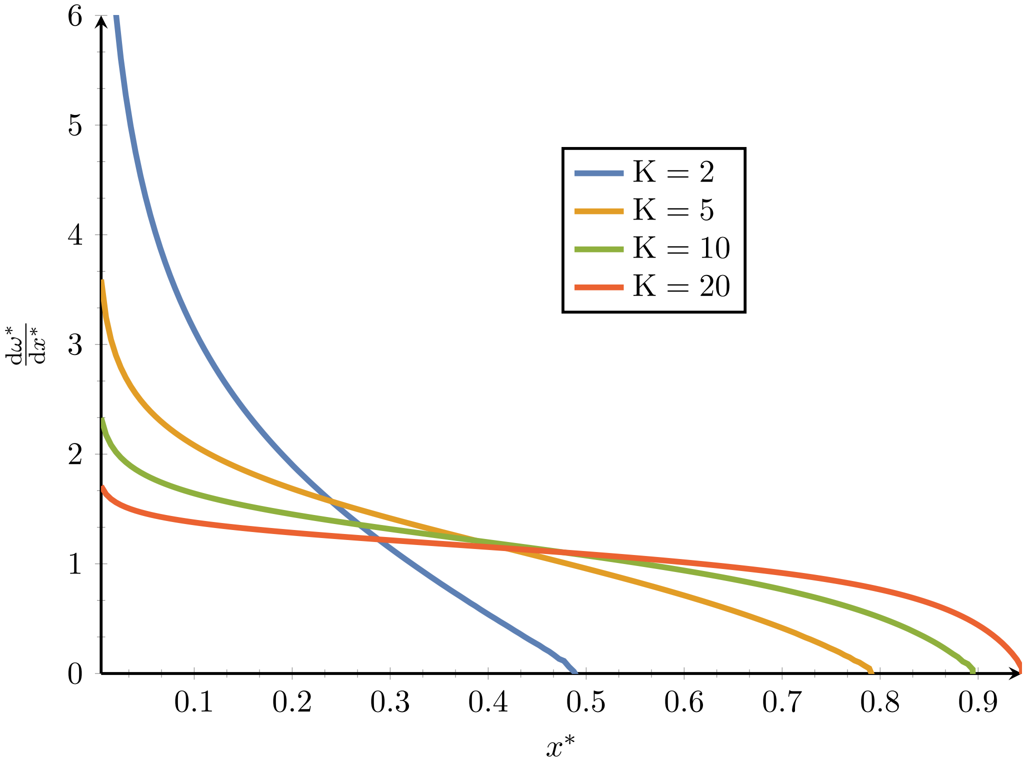

Moreover, given that the number of state if large enough $\mathrm{K} \gg 1$, the response in equilibrium $\omega$ due to change in phenotype can be approximated as:

$$\frac{\mathrm{d}\omega^*}{\mathrm{d}x^*} = 2\left[\frac{\mathrm{K}(x^*-1)+1+\left[\mathrm{K}(x^*-1)^2+2x^*-1\right]\lambda_K(x^*)}{(\mathrm{K}(x^*-1)+1)^2}\right] + \frac{\mathrm{K}-2}{\mathrm{K}-1}, \tag{28}$$

$$\simeq \frac{2\lambda_K(x^*)}{\mathrm{K}} + 1, \tag{29}$$

$$\simeq 1. \tag{30}$$

#### 1.6 $\omega$ response after a change in $N_\mathrm{e}$

Define the function $G(x, N_\mathrm{e})$ as:

$$G(x, N_\mathrm{e}) \equiv \lambda_K(x^*) + 4N_\mathrm{e} s(x, \delta x), \tag{31}$$

The equilibrium equation (eq. 18) states that $G(x^*, N_\mathrm{e}) = 0$, meaning that $x^*$ is implicitly a function of $N_\mathrm{e}$:

$$G(x^*(N_\mathrm{e}), N_\mathrm{e}) = 0, \tag{32}$$

$$\implies \frac{\partial G(x^*, N_\mathrm{e})}{\partial x^*}\frac{\mathrm{d}x^*}{\mathrm{d}N_\mathrm{e}} + \frac{\partial G(x^*, N_\mathrm{e})}{\partial N_\mathrm{e}} = 0, \tag{33}$$

$$\iff \left[\frac{\partial \lambda_K(x^*)}{\partial x^*} + 4N_\mathrm{e}\frac{\partial s(x^*, \delta x)}{\partial x^*}\right]\frac{\mathrm{d}x^*}{\mathrm{d}N_\mathrm{e}} + 4s(x^*, \delta x) = 0, \tag{34}$$

$$\iff \left[\frac{\partial \lambda_K(x^*)}{\partial x^*} + 4N_\mathrm{e}\frac{\partial^2 f(x^*)}{\partial x^{*2}}\delta x\right]\frac{\mathrm{d}x^*}{\mathrm{d}N_\mathrm{e}} = -4\frac{\partial f(x^*)}{\partial x^*}\delta x \text{ from eq. 5}, \tag{35}$$

$$\iff 4\delta x\left[\frac{1}{4\delta x N_\mathrm{e}}\frac{\partial \lambda_K(x^*)}{\partial x^*} + \frac{\partial^2 f(x^*)}{\partial x^{*2}}\right] N_\mathrm{e}\frac{\mathrm{d}x^*}{\mathrm{d}N_\mathrm{e}} = -4\delta x\frac{\partial f(x^*)}{\partial x^*}, \tag{36}$$

$$\iff \frac{\mathrm{d}x^*}{\mathrm{d}\ln(N_\mathrm{e})} = -\frac{\frac{\partial f(x^*)}{\partial x^*}}{\frac{1}{4\delta x N_\mathrm{e}}\frac{\partial \lambda_K(x^*)}{\partial x^*} + \frac{\partial^2 f(x^*)}{\partial x^{*2}}}. \tag{37}$$

Giving the equation for the response of phenotype at equilibrium after a change of effective population size. Together, the response of substitution rate at equilibrium, after a change of effective population size can be obtained as:

$$\frac{\mathrm{d}\omega^*}{\mathrm{d}\ln(N_\mathrm{e})} = \frac{\mathrm{d}\omega^*}{\mathrm{d}x^*}\frac{\mathrm{d}x^*}{\mathrm{d}\ln(N_\mathrm{e})}, \tag{38}$$

$$= -\frac{\mathrm{d}\omega^*}{\mathrm{d}x^*}\frac{\frac{\partial f(x^*)}{\partial x^*}}{\frac{1}{4\delta x N_\mathrm{e}}\frac{\partial \lambda_K(x^*)}{\partial x^*} + \frac{\partial^2 f(x^*)}{\partial x^{*2}}} \text{ from eq. 37.} \tag{39}$$

Moreover, with the approximation that $\left|4N_\mathrm{e}\frac{\partial s(x^*,\delta x)}{\partial x^*}\right| \gg \left|\frac{\partial \lambda_K(x^*)}{\partial x^*}\right|$, meaning that a change in phenotype causes a higher change in scaled selection coefficient than mutational bias, we have:

$$\frac{\mathrm{d}x^*}{\mathrm{d}\ln(N_\mathrm{e})} = -\frac{\frac{\partial f(x^*)}{\partial x^*}}{\frac{1}{4\delta x N_\mathrm{e}}\frac{\partial \lambda_K(x^*)}{\partial x^*} + \frac{\partial^2 f(x^*)}{\partial x^{*2}}}, \tag{40}$$

$$\implies \frac{\mathrm{d}x^*}{\mathrm{d}\ln(N_\mathrm{e})} \simeq -\frac{\frac{\partial f(x^*)}{\partial x^*}}{\frac{\partial^2 f(x^*)}{\partial x^{*2}}}. \tag{41}$$

Together, these approximations leads to the following response in equilibrium $\omega$ after change in $N_\mathrm{e}$ as:

$$\frac{\mathrm{d}\omega^*}{\mathrm{d}\ln(N_\mathrm{e})} \simeq -\frac{\frac{\partial f(x^*)}{\partial x^*}}{\frac{\partial^2 f(x^*)}{\partial x^{*2}}} \tag{42}$$

### 2 Models for the log-fitness function

#### 2.1 Folded fraction

All phenotype-fitness functions considered below are log-concave, and as a result, $\frac{\partial f(x^*)}{\partial x^*}$ is a decreasing function of $x$; the less stable the protein already is, the stronger the purifying selection against additional destabilizing mutations. More precisely, fitness functions depends on the folded fraction of the protein of interest, which is given by the Fermi-Dirac distribution:

$$\mathbb{P}_\mathrm{F}(x) = \frac{1}{1+e^{\beta(\alpha+\gamma n x)}}, \tag{43}$$

where $x$ is the fraction of destabilizing mutations, each contributing to $\gamma$ in free energy of folding (also denoted empirically $\Delta\Delta G$), and $\beta = 1/kT$. Thus, $\alpha < 0$ is the difference in free energy between folded and unfolded state when all sites are stable (also denoted empirically $\Delta G_\mathrm{min}$). As a result, $n\gamma$ is thus the expected change in $\Delta G$ when all sites are considered unstable. The misfolded fraction is $\mathbb{P}_\mathrm{U} = 1 - \mathbb{P}_\mathrm{F}$. In addition, $\mathbb{P}_\mathrm{F}$ is typically close to 1 (or $\mathbb{P}_\mathrm{U} \ll 1$), so that we can use a first-order approximation:

$$\mathbb{P}_\mathrm{F}(x) = 1 - \mathbb{P}_\mathrm{U}(x) \tag{44}$$

$$\simeq 1 - e^{\beta(\alpha+\gamma n x)} \tag{45}$$

or equivalently

$$\mathbb{P}_\mathrm{U}(x) \simeq e^{\beta(\alpha+\gamma n x)} \tag{46}$$

#### 2.2 Fitness equal to folded fraction

A first model is to assume that the fitness is equal to the folded fraction (Goldstein, 2013):

$$W(x) = \frac{1}{1+\mathrm{e}^{\beta(\alpha+n\gamma x)}}. \tag{47}$$

The derivative of fitness w.r.t to phenotype is:

$$\frac{\partial f(x)}{\partial x} = -\frac{\partial \ln\left(1+\mathrm{e}^{\beta(\alpha+n\gamma x)}\right)}{\partial x} \text{ from eq. 47,} \tag{48}$$

$$= -\beta n\gamma \frac{\mathrm{e}^{\beta(\alpha+n\gamma x)}}{1+\mathrm{e}^{\beta(\alpha+n\gamma x)}}, \tag{49}$$

$$\simeq -\beta n\gamma \mathrm{e}^{\beta(\alpha+n\gamma x)}. \tag{50}$$

The equilibrium phenotype ($x^*$) is :

$$\lambda_K(x^*) = 4N_\mathrm{e}\beta\gamma \frac{\mathrm{e}^{\beta(\alpha+n\gamma x^*)}}{1+\mathrm{e}^{\beta(\alpha+n\gamma x^*)}} \text{ from eq. 19 and 49.} \tag{51}$$

Using $N_\mathrm{e} = 10^4$, $\beta = 1.686$, $\alpha = \Delta G_\mathrm{min} = -118$, $n = 300$, $\gamma = \Delta\Delta G = 1$, we have the following :

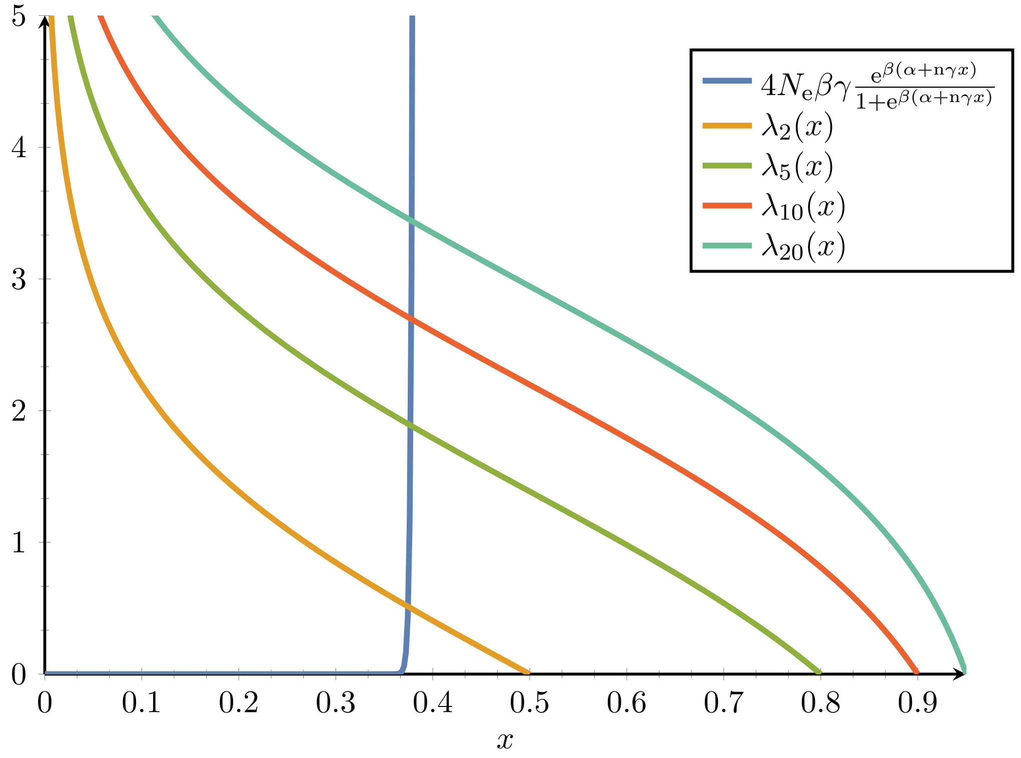

Where in this example we can visually appreciate that the a change in phenotype causes a higher change in scaled selection coefficient than mutational bias (eq. 41). And the second derivative of fitness w.r.t to phenotype is:

$$\frac{\partial^2 f(x)}{\partial x^2} = -\beta n\gamma \frac{\partial}{\partial x}\left(\frac{\mathrm{e}^{\beta(\alpha+n\gamma x)}}{1+\mathrm{e}^{\beta(\alpha+n\gamma x)}}\right) \text{ from eq. 49,} \tag{52}$$

$$= -\beta n\gamma\beta n\gamma \frac{\mathrm{e}^{\beta(\alpha+n\gamma x)}}{\left(1+\mathrm{e}^{\beta(\alpha+n\gamma x)}\right)^2}, \tag{53}$$

$$= \frac{\beta n\gamma}{1+\mathrm{e}^{\beta(\alpha+n\gamma x)}} \frac{\partial f(x)}{\partial x} \text{ from eq. 49,} \tag{54}$$

$$\simeq \beta n\gamma \frac{\partial f(x)}{\partial x} \tag{55}$$

Finally, $\omega$ response after a change in $N_\mathrm{e}$ is simply:

$$\frac{\mathrm{d}\omega^*}{\mathrm{d}\ln(N_\mathrm{e})} \simeq -\frac{1}{\beta n\gamma} \text{from eq. 54 and 30,} \tag{56}$$

which is independent of $x^*$, meaning $\omega$ is linearly decreasing with $N_\mathrm{e}$ in log space. This model, however, does not express the fact that selection is typically stronger for proteins characterized by higher levels of expression.

#### 2.3 Selective cost proportional to amount of misfolded protein

A slight variation is to assume that the selective cost itself is proportional to the total amount of misfolded protein (Drummond *et al.*, 2005; Wilke and Drummond, 2006; Drummond and Wilke, 2008; Serohijos *et al.*, 2012). For a given protein with expression level $y$:

$$f(x) = -Ay\mathbb{P}_{\mathrm{U}}(x), \tag{57}$$

where $A$ is the cost per misfolded macromolecule. Then,

$$\frac{\partial f(x)}{\partial x} \simeq -Ay\beta\gamma n e^{\beta(\alpha+\gamma n x)}. \tag{58}$$

Under this model, the phenotype at equilibrium is given by:

$$\lambda_K(x^*) = 4N_{\mathrm{e}}yA\beta\gamma n e^{\beta(\alpha+\gamma n x^*)} \text{ from eq. 19 and 49.} \tag{59}$$

And the response of $\omega$ after a change in $N_{\mathrm{e}}$ is the same as before:

$$\frac{\mathrm{d}\omega^*}{\mathrm{d}\ln(N_{\mathrm{e}})} \simeq -\frac{1}{\beta n\gamma}. \tag{60}$$

Since $N_{\mathrm{e}}$ and $y$ are confounded factors, meaning they only appear in the equation as a product between the two, implicit derivation leads to the same result whenever the derivation is w.r.t $N_{\mathrm{e}}$ or $y$, leading to same compact equation:

$$\frac{\mathrm{d}\omega^*}{\mathrm{d}\ln(y)} = \frac{\mathrm{d}\omega^*}{\mathrm{d}\ln(N_{\mathrm{e}})} \simeq -\frac{1}{\beta n\gamma}. \tag{61}$$

#### 2.4 Translational errors

Another variant account for translational errors. Translational errors occur at a rate $\rho$ per residue. These errors contribute additional destabilizing mutations, each with effect size $\delta x = 1/n$. The total number of translational errors per macromolecule is approximately Poisson distributed:

$$\pi_k = e^{-\rho n}\frac{(\rho n)^k}{k!} \tag{62}$$

and the total selective cost is now an average over all possible values of $k$:

$$\begin{aligned} f(x) &= -Ay\sum_k \pi_k e^{\beta(\alpha+\gamma n x+\gamma k)} && (63)\\ &= -Aye^{\beta(\alpha+\gamma n x)}\sum_k e^{-\rho n}\frac{(\rho n)^k}{k!}e^{\beta\gamma k} && (64)\\ &= -Aye^{\beta(\alpha+\gamma n x)+\rho n(e^{\beta\gamma}-1)} && (65)\\ &\simeq -Aye^{\beta(\alpha+\gamma n x)+\rho\beta\gamma n} && (66)\\ &= -Aye^{\beta(\alpha+\gamma n(x+\rho))} && (67)\end{aligned}$$

In words, the fitness function is the same the previous model, except that the trait $x$ (fraction of destabilizing mutations) is shifted by $\rho$, the mean fraction of additional mutations contributed by translation errors. This additional factor is independent of $x$, and as a result, the scaled selection strength is essentially the same, up to a proportionality constant (contributed by the shift):

$$\begin{aligned} 4N_{\mathrm{e}}\frac{\partial f(x)}{\partial x} &\propto -4N_{\mathrm{e}}ye^{\beta(\alpha+\gamma n(x+\rho))} && (68)\\ &\propto -4N_{\mathrm{e}}ye^{\beta(\alpha+\gamma n x)} && (69)\end{aligned}$$

Moreover, $\omega$ response after a change in $N_{\mathrm{e}}$ is again the same as before:

$$\frac{\mathrm{d}\omega^*}{\mathrm{d}\ln(N_{\mathrm{e}})} \simeq -\frac{1}{\beta n\gamma}. \tag{70}$$

##### 2.5 Cost-benefit argument

The cost-benefit argument (Beaulieu *et al.*, 2018) is based on two assumptions

1. the expression level is regulated so that the total number of *functional* macromolecules is maintained at a target level $y$;
2. the log-fitness is proportional to the ratio of the *total* cost of expression over the benefit contributed by the protein.

Specifically, the protein is assumed to be regulated so as to reach a level of expression of functional proteins of $y$, and contributes a total benefit $B$ (which depends on its specific function). Given that only a fraction $\mathbb{P}_{\mathrm{F}}(x) = 1 - \mathbb{P}_{\mathrm{U}}(x)$ of the total amount of protein expressed by the cell is functional, the total cost of expression $C$ is then equal to:

$$C(x) = \frac{y}{\mathbb{P}_{\mathrm{F}}(x)} \tag{71}$$

$$\simeq y(1 + \mathbb{P}_{\mathrm{U}}(x)) \tag{72}$$

Then, the log-fitness is given by:

$$f(x) = -A\frac{y}{B}\left(1 + e^{\beta(\alpha+\gamma n x)}\right) \tag{73}$$

$$= -by(1 + e^{\beta(\alpha+\gamma n x)}), \tag{74}$$

where $b = A/B$. Compared to models 2 (section 2.3) and 3 (section 2.4), the log-fitness now has an additional term that depends on the target expression level $y$, but not on trait $x$. The scaled strength of selection on mutations affecting $x$ has thus the same functional form as for the two previous models:

$$4N_{\mathrm{e}}\frac{\partial f(x)}{\partial x} \propto -4N_{\mathrm{e}}ye^{\beta(\alpha+\gamma n x)} \tag{75}$$

Alternative cost-expression models could also be used, allowing for a non-linear cost function for expression or for some susceptibility of the realized equilibrium expression level, as a function of the number of mutations. Under these models, the strength of selection is still expected to be an increasing function of $y$, although not linear:

$$4N_{\mathrm{e}}\frac{\partial f(x)}{\partial x} \propto -4N_{\mathrm{e}}g(y)e^{\beta(\alpha+\gamma n x)}, \tag{76}$$

where $g$ is some function of $y$. Moreover, $\omega$ response after a change in $N_{\mathrm{e}}$ is again the same as before:

$$\frac{\mathrm{d}\omega^*}{\mathrm{d}\ln(N_{\mathrm{e}})} \simeq -\frac{1}{\beta n\gamma}. \tag{77}$$

#### 3 Model of protein-protein interactions

The proteome is assumed to be composed of $m$ protein species, all with same abundance $C$. Each macromolecule may either be in free form or engaged in a non-specific interaction. Only pairwise interactions are considered, and higher-order interactions are ignored. The equilibrium is characterized by:

$$[ij] = \frac{[i][j]}{C_0}\, e^{\beta E_{ij}}, \tag{78}$$

where $[i]$ and $[j]$ are the concentrations of protein species $i$ and $j$, and $[ij]$ is the concentration of their (non-specific) dimer. Here, $E_{ij}$ is the interaction free energy, which can itself be decomposed as a sum of three terms:

$$E_{ij} = \alpha + E_i + E_j \tag{79}$$

$$= \alpha + \gamma n(x_i + x_j), \tag{80}$$

where we assume that each protein has $n = 100$ residues at its surface, $x_i$ stands for the fraction of hydrophobic residues at the surface of protein $i$, and each hydrophobic residue makes an additive contribution of $\Delta\Delta G$ to the total.

By conservation of the total number of molecules:

$$C = [i] + \sum_{j \neq i} [ij] \tag{81}$$

$$= [i] + \sum_{j \neq i} \frac{[i][j]}{C_0} \, e^{\beta E_{ij}} \tag{82}$$

and we note:

$$\epsilon_i = \sum_j [ij] \tag{83}$$

the fraction of protein $i$ sequestered in non-specific interactions. We assume that the log fitness is proportional to the total amount of protein sequestered in non-specific interactions:

$$f(x) = -b \sum_i \epsilon_i, \tag{84}$$

where $b > 0$ is a parameter determining the overall stringency of selection against non-specific interactions.

#### 3.1 Mean field, weak-interaction limit

To make the model tractable and compact, we assume that non-specific interactions are weak, i.e. $\epsilon_i \ll 1$ for all $i$. We then make a first-order approximation in the $\epsilon_i$'s. In addition, we use a mean-field approximation, such that, when considering a specific protein species $i$, we assume that all other proteins have the same fraction $\bar{x}$ of hydrophobic residues at their surface. The value of $\bar{x}$ could in principle be found using a self-consistent argument, essentially by (1) explicitly calculating the net substitution flux for protein $i$ with fraction $x_i$, under mean field $\bar{x}$, and (2) expressing the constraint that this substitution process for protein $i$ is stationary at $x_i = \bar{x}$. This derivation is not conducted here, as it is not needed. Using these approximations, we can re-express the conservation of total mass as:

$$C = [i] + (m-1)[i] \frac{C}{C_0} \, e^{\beta(\alpha + \gamma n(\bar{x} + x_i))} \tag{85}$$

Here, we have used the fact that $[j] = C(1 - \epsilon_j)$ can be approximated as $[j] \simeq C$ since it is involved in a term already of the order of $\epsilon_i$. As a result, all $m-1$ terms of the sum over $j \neq i$ are identical. Next, solving for $[i]$ gives:

$$[i] = \frac{C}{1 + (m-1)\frac{C}{C_0} \, e^{\beta(\alpha + \gamma n(\bar{x} + x_i))}} \tag{86}$$

$$\simeq C \left( 1 - m \frac{C}{C_0} \, e^{\beta(\alpha + \gamma n(\bar{x} + x_i))} \right) \tag{87}$$

$$= C(1 - \epsilon_i) \tag{88}$$

and thus $\epsilon_i$ can be identified with:

$$\epsilon_i = m \frac{C}{C_0} \, e^{\beta(\alpha + \gamma n(\bar{x} + x_i))} \tag{89}$$

Now, assume that the system is at equilibrium (thus $x_i = \bar{x}$). The strength of selection acting on mutations occurring at the surface of protein $i$, of effect size $\delta x = \pm 1/n$, is given by $s = \kappa \delta x$ where:

$$\kappa_i = b \frac{\mathrm{d}\epsilon_i}{\mathrm{d}x_i} \tag{90}$$

$$= b\beta\gamma n m \frac{C}{C_0} \, e^{\beta(\alpha + \gamma n(\bar{x} + x_i))} \tag{91}$$

and thus:

$$\ln(\kappa_i) = \ln\left(b\beta\gamma nm\frac{C}{C_0}\right) + \beta(\alpha + \gamma n\bar{x}) + \beta\gamma n x_i, \tag{92}$$

where only the last term depends on $x_i$. Finally, applying the main result of this work to the present case allows us to express the response of $\omega$ as a function of $N_\mathrm{e}$ as:

$$\chi = \frac{\mathrm{d}\omega}{\mathrm{d}\ln(N_\mathrm{e})} \tag{93}$$

$$= 2(\lambda - 1)\,\frac{\mathrm{d}\ln\kappa_i}{\mathrm{d}x_i} \tag{94}$$

$$= 2(\lambda - 1)\,\frac{1}{\beta\gamma n} \tag{95}$$

Note that, here, we have used $K = 2$ (hydrophobic and polar residues are roughly equally likely to occur by mutation), and assumed $x^* \ll 1$. A more accurate formula could be used without this latter assumption. In any case, $\chi$ is now dependent on $x^*$, through $\lambda$.

##### 3.2 Empirical calibration

Based on empirical estimates found in Zhang *et al.* (2008). The mean fraction of hydrophobic residues at the surface of proteins is $0.22 \pm 0.06$. With $n = 100$ residues, this makes $22 \pm 6$. The mean value for $E_{ij}$ is $7kT$, with a standard deviation of $\sigma = 1.8kT$. Assuming that this standard deviation of $\pm 1.8kT$ is contributed by $\pm 6$ mutations gives $\Delta\Delta G = 1.8/6 = 0.3$ kT or 0.18 kcal per mole. Also, with $x = 0.22$, $\lambda \simeq 4$, and thus $\chi = 6/30 = 0.2$, thus a much stronger response than under the model based on conformational stability.

#### 4 Empirical estimation

| **Type** | **Specie** | $\widehat{\chi}$ | $r^2$ |
|---|---|---|---|
| Plant | Oryza sativa | -0.008 | 0.047 |
| Plant | Arabidopsis thaliana | -0.012 | 0.128 |
| Archaea | Sulfolobus solfataricus | -0.037 | 0.097 |
| Archaea | Thermococcus kodakarensis | -0.026 | 0.058 |
| Fungi | Saccharomyces cerevisiae | -0.029 | 0.211 |
| Fungi | Aspergillus nidulans | -0.034 | 0.124 |
| Bacteria | Escherichia coli | -0.021 | 0.151 |
| Bacteria | Bacillus subtilis | -0.046 | 0.151 |
| Animal | Caenorhabditis elegans | -0.026 | 0.039 |
| Animal | Drosophila melanogaster | -0.005 | 0.021 |
| Animal | Mus musculus | -0.008 | 0.085 |
| Animal | Homo sapiens | -0.004 | 0.031 |

Table 1: Substitution rate as a function of expression level compiled by Zhang and Yang (2015).

In Brevet and Lartillot (2019), the covariance matrix with $\ln(N_\mathrm{e})$ and $\ln(\omega)$ as entries allows to approximate $\chi$:

$$\widehat{\frac{\mathrm{d}\ln(\omega)}{\mathrm{d}\ln(N_\mathrm{e})}} = \frac{\mathrm{Cov}[\ln(\omega), \ln(N_\mathrm{e})]}{\mathrm{Var}[\ln(N_\mathrm{e})]}, \tag{96}$$

$$\Rightarrow \widehat{\frac{\mathrm{d}\omega}{\omega\mathrm{d}\ln(N_\mathrm{e})}} = \frac{\mathrm{Cov}[\ln(\omega), \ln(N_\mathrm{e})]}{\mathrm{Var}[\ln(N_\mathrm{e})]}, \tag{97}$$

$$\Rightarrow \widehat{\chi} \simeq \widehat{\omega}\frac{\mathrm{Cov}[\ln(\omega), \ln(N_\mathrm{e})]}{\mathrm{Var}[\ln(N_\mathrm{e})]}, \tag{98}$$

$$\Rightarrow \widehat{\chi} \simeq 0.2\frac{-0.45}{4.45}, \tag{99}$$

$$\Rightarrow \widehat{\chi} \simeq -0.02 \tag{100}$$

### 5 Simulation using the 3D structure of protein

We simulated substitutions in the protein phosphatase ($\mathrm{Z} = 300$ codon sites) as in Goldstein and Pollock (2017). From a DNA sequence $\mathbb{S}$ after $t$ substitutions, we compute the free energy of the folded state $G_{\mathrm{F}}(\mathbb{S})$, using the 3-dimensional structure of the folded state and pair-wise contact energies between neighboring amino-acid residues:

$$G_{\mathrm{F}}(\mathbb{S}) = \sum_{\mathrm{z}=1}^{\mathrm{Z}} \sum_{r \in \mathcal{V}(\mathrm{z})} I(\mathbb{S}(\mathrm{z}), \mathbb{S}(r)), \tag{101}$$

where $I(a, b)$ is the pair-wise contact energies between amino acid $a$ and $b$, using contact potentials estimated by Miyazawa and Jernigan (1985), and $\mathcal{V}(\mathrm{z})$ are the neighbor residues of site z (closer than 7Å) in the 3D structure.

The free energy of unfolded states $G_{\mathrm{U}}(\mathbb{S})$ is approximated using 55 decoy 3D structures that supposedly represent a sample of possible unfolded states:

$$G_{\mathrm{U}}(\mathbb{S}) = \langle G(\mathbb{S}) \rangle - kT \ln(1.0\mathrm{E}^{160}) - \frac{2\left[\left\langle G(\mathbb{S})^2 \right\rangle - \langle G(\mathbb{S}) \rangle^2\right]}{kT} \tag{102}$$

where the average $\langle . \rangle$ runs other the 55 decoy 3D structures, and $k$ is the Boltzmann constant and $T$ the temperature in Kelvin.

From the energy of folded and unfolded states, we can compute the difference in free energy between the states:

$$\Delta G(\mathbb{S}) = G_{\mathrm{F}}(\mathbb{S}) - G_{\mathrm{U}}(\mathbb{S}) \tag{103}$$

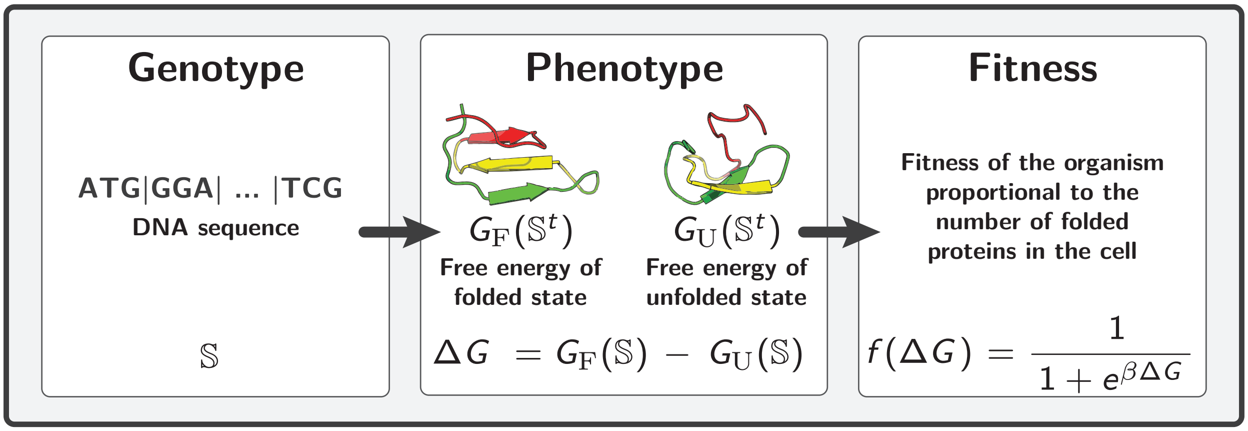

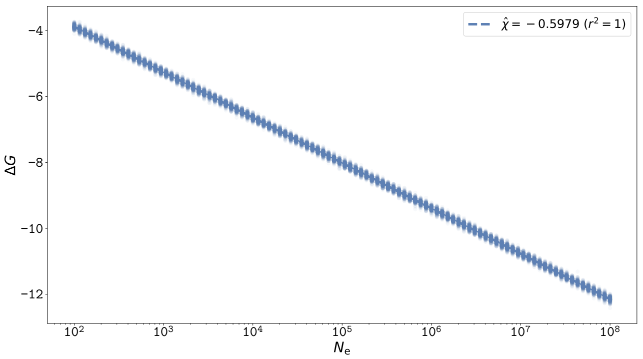

Figure 1: $\Delta G$ response to changes in $N_{\rm e}$. Left Panel: Model of folding free energy computed using 3D structural conformations and pairwise contact potential energies between neighbouring amino-acid residues. Right Panel: Additive phenotype model, where for each non-optimal amino acid, $\gamma$ is scaled by the Grantham distance to the optimal amino acid. Scaling experiment simulating sequence evolution and recording the average $\Delta G$ (y-axis) observed at equilibrium as a function of $N_{\rm e}$ (x-axis). Along the x-axis, 200 replicate simulations are performed for each different $N_{\rm e}$, the average (solid lines) and 90% confidence interval (shaded area) of $\omega$ are shown. $\Delta G$ is linearly dependent on log-$N_{\rm e}$, with a slope equal to $1/\beta = 0.593$.

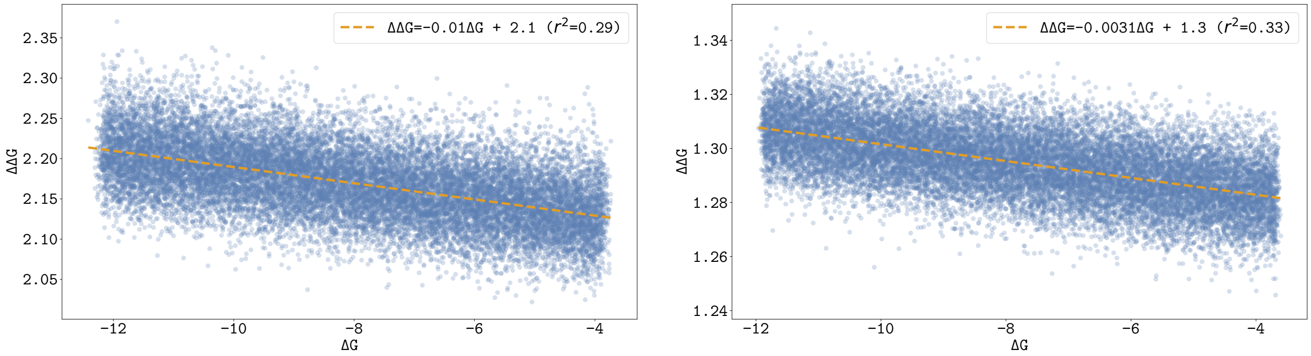

Figure 2: $\Delta\Delta G$ correlation to $\Delta G$. Left Panel: Model of folding free energy computed using 3D structural conformations and pairwise contact potential energies between neighbouring amino-acid residues. Right Panel: Additive phenotype model, where for each non-optimal amino acid, $\gamma$ is scaled by the Grantham distance to the optimal amino acid. Simulations are performed for $N_{\rm e}$ varying from $10^2$ to $10^8$, where each dot is an independent simulation at equilibrium. Along each simulation, the average $\Delta\Delta G$ of all proposed mutations is recorded (y-axis), and represented as a function of the average $\Delta G$ (x-axis). $\Delta\Delta G$ is negatively correlated to $\Delta$, which is expected since protein under higher $N_{\rm e}$ are more stable (lower $\Delta G$, see above), and mutations are more destabilizing on average. To be more precise, the negative correlation between $\Delta\Delta G$ and $\Delta G$ is observed empirically with a linear fit of $\Delta\Delta G = -0.13\Delta G + 0.23$ (Serohijos *et al.*, 2012). This correlation is a necessary condition for observing a response of $\omega$ to changes in $N_{\rm e}$ (Goldstein, 2013) and protein expression level (Serohijos *et al.*, 2012).

### 6 Simulated $\omega$ response to changes in $N_e$

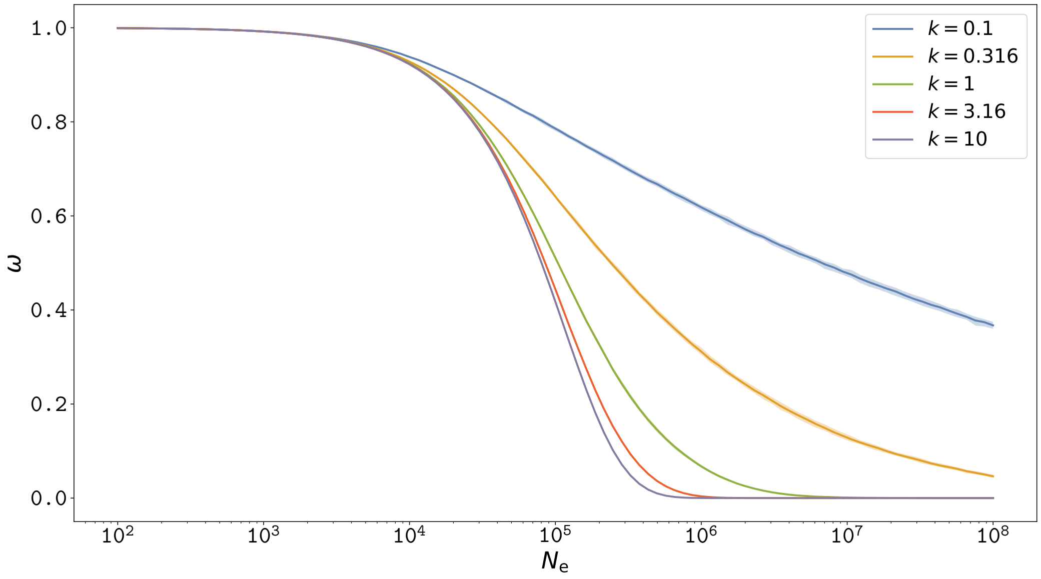

Figure 3: $\omega$ at equilibrium as a function of $N_e$ (log scale), under a model of gamma distributed selection coefficient. For each population size, 200 simulations were performed and the average (solid line) and 90% confidence interval (shaded area) are shown. In the model of gamma distributed fitness effect, $\omega$ at equilibrium is strongly dependent on log-$N_e$ where the slope correlation is proportional to the inverse of the shape parameter of the gamma distribution (Welch *et al.*, 2008).

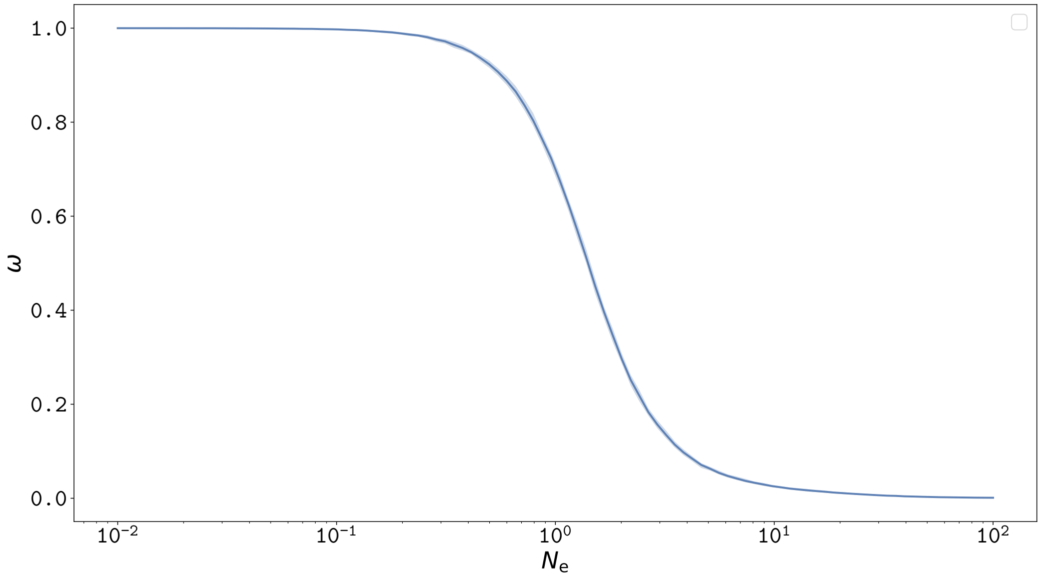

Figure 4: $\omega$ at equilibrium as a function of $N_e$ (relative), under a model of amino-acid fitness profiles. For each population size, 200 simulations were performed and the average (solid line) and 90% confidence interval (shaded area) are shown. In the model of site-wise amino-acid fitness profiles taken from (Bloom, 2017), $\omega$ at equilibrium is strongly dependent on log-$N_e$.

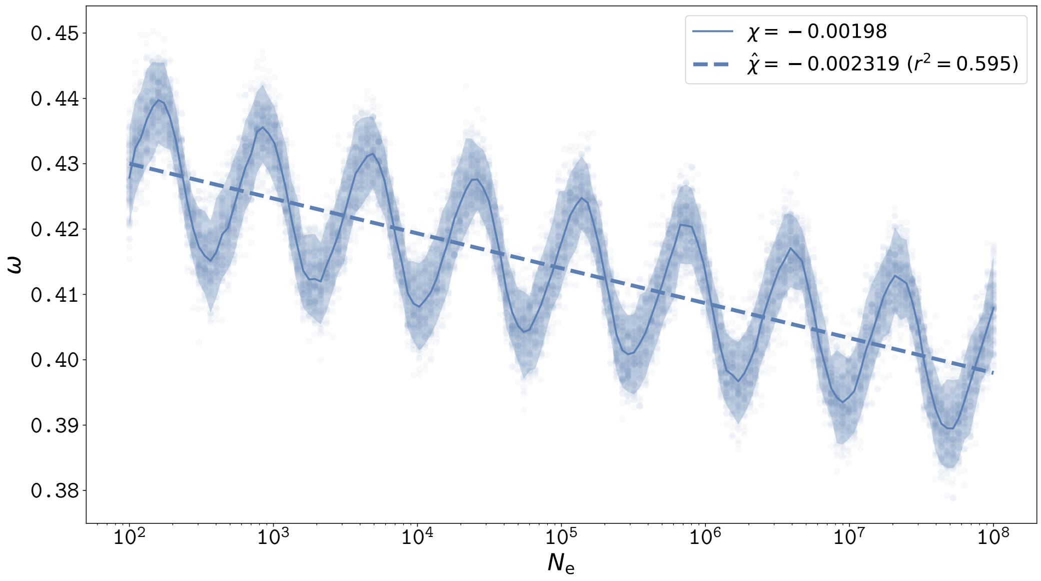

Figure 5: $\omega$ at equilibrium as a function of $N_e$ (log scale), for a model of additive free energy of folding. For each population size, 200 simulations were performed and the average (solid line) and 90% confidence interval (shaded area) are shown. The fixed parameters are $\alpha = -118$, $\gamma = 1$, $n = 300$, $\beta = 1.686$. The simulations of our additive free energy model match the theoretical prediction that the slope of the linear relation (dashed line) is equal to $\beta n\gamma)^{-1} = 0.00198 \simeq= 0.00199$. The non-monotony is suspected to be due to the discrete number of sites and states, such that the changes in $\Delta G$ after a mutation is either $-1$, 0 or 1. Such non-monotony is not observed with the Grantham model, in which the $\omega$ is lower and the slope of the response is lower, closer to the empirical 3D model.

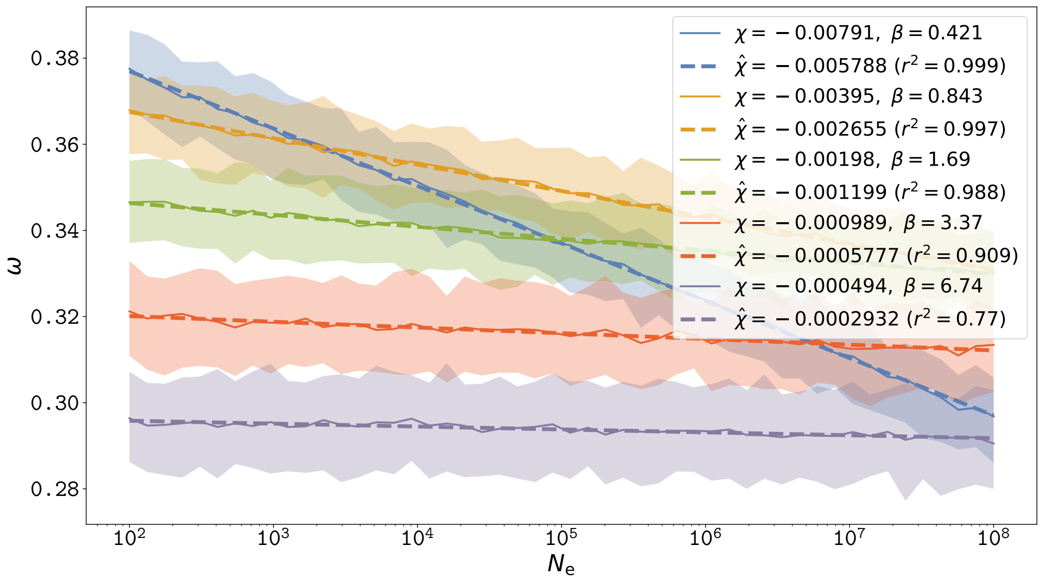

Figure 6: $\omega$ at equilibrium as a function of $N_e$ (log scale), for various parameter $\beta$. For each population size, 200 simulations were performed and the average (solid line) and 90% confidence interval (shaded area) are shown. The fixed parameters are $\alpha = -118$, $\gamma = 1$, $n = 300$, and for each non-optimal amino acid, $\gamma$ is scaled by the Grantham distance to the optimal amino acid. $\beta$ are given in the legend. Increasing $\beta$ decreases the slope of the $\omega$-$N_e$ relationship, as predicted in our theoretical model.

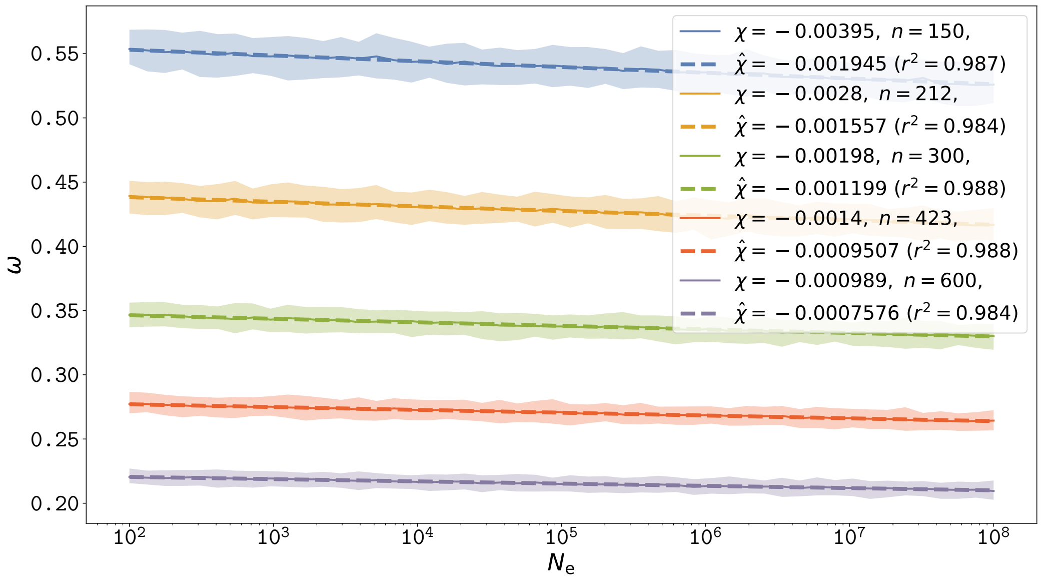

Figure 7: $\omega$ at equilibrium as a function of $N_e$ (log scale), for various sequence size. For each population size, 200 simulations were performed and the average (solid line) and 90% confidence interval (shaded area) are shown. The fixed parameters are $\alpha = -118$, $\gamma = 1$, $\beta = 1.686$, and for each non-optimal amino acid, $\gamma$ is scaled by the Grantham distance to the optimal amino acid. $n$ are given in the legend. Increasing $n$ decreases the slope of the $\omega$-$N_e$ relationship, as predicted in our theoretical model.

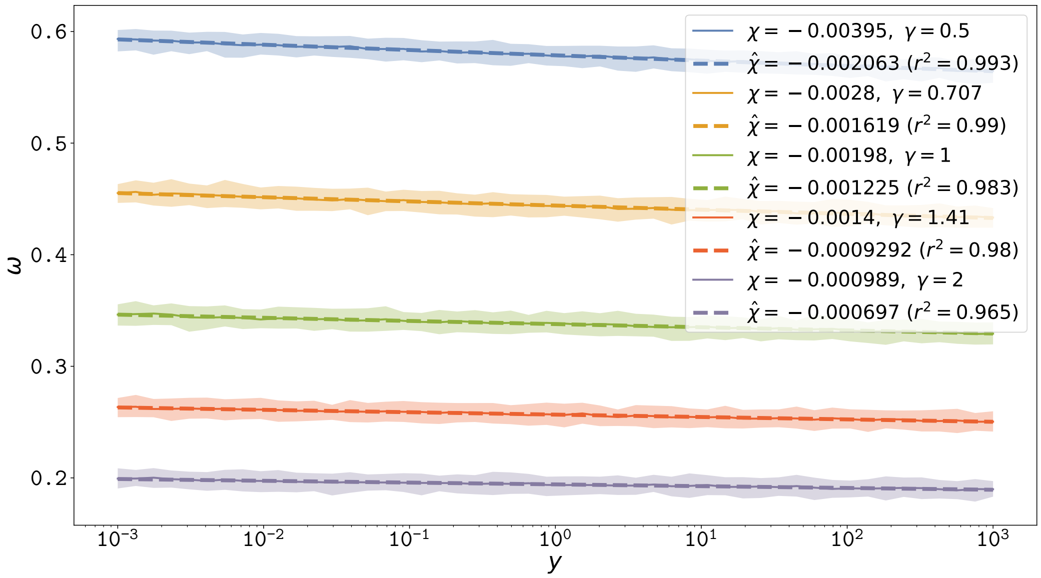

Figure 8: $\omega$ at equilibrium as a function of the expression level $y$ (log scale), for various value of $\gamma$. For each population size, 200 simulations were performed and the average (solid line) and 90% confidence interval (shaded area) are shown. The fixed parameters are $\alpha = -118$, $\beta = 1.686$, $n = 300$, and for each non-optimal amino acid, $\gamma$ is scaled by the Grantham distance to the optimal amino acid. $\gamma$ are given in the legend. Increasing $\gamma$ increases the slope of the $\omega$-$y$ relationship, as predicted in our theoretical model.

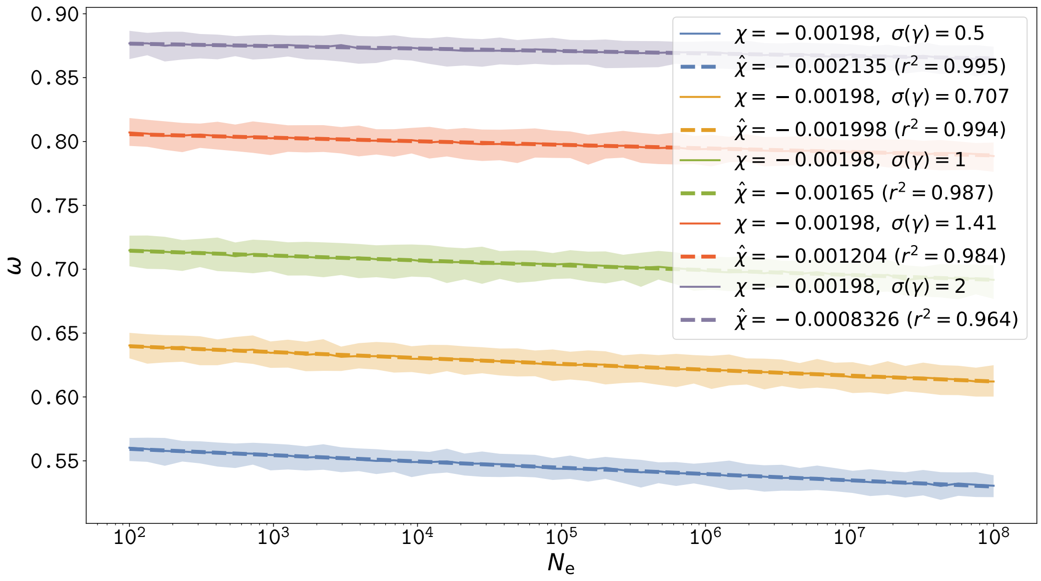

Figure 9: $\omega$ at equilibrium as a function of $N_e$ (log scale), for various between site variance. For each population size, 200 simulations were performed and the average (solid line) and 90% confidence interval (shaded area) are shown. The parameters are $\alpha = -118$, $n = 300$, $\beta = 1.686$ and each site has its own gamma distributed $\gamma$ with mean 1 and standard deviation given in the legend. $\gamma$ is scaled by the Grantham distance to the optimal amino acid. Increasing the variance of $\gamma$ increases $\omega$, by shifting the equilibrium to higher $x^*$ since more unstable sites with low $\gamma$ are fixed before reaching sensible deleterious selection coefficient against unstable mutations. Once many sites are unstable, the $\omega$ is higher since non-synonymous mutations between unstable states are effectively neutral. However the slope of the $\omega$-$N_e$ relationship is not sensibly changed.

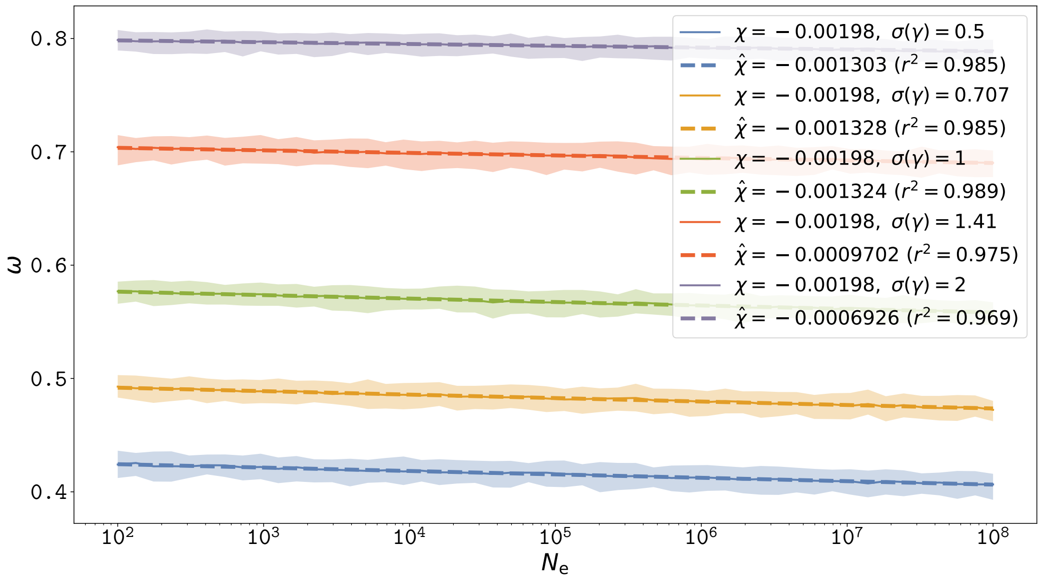

Figure 10: $\omega$ at equilibrium as a function of $N_\mathrm{e}$ (log scale), for various between site variance under a model considering Grantham distances. For each population size, 200 simulations were performed and the average (solid line) and 90% confidence interval (shaded area) are shown. The parameters are $\alpha = -118$, $n = 300$, $\beta = 1.686$ and each site has its own gamma distributed $\gamma$ with mean 1 and standard deviation given in the legend. Increasing the variance of $\gamma$ increases $\omega$, by shifting the equilibrium to higher $x^*$ since more unstable sites with low $\gamma$ are fixed before reaching sensible deleterious selection coefficient against unstable mutations. Once many sites are unstable, the $\omega$ is higher since non-synonymous mutations between unstable states are effectively neutral. However the slope of the $\omega$-$N_\mathrm{e}$ relationship is not sensibly changed.

### 7 Simulated relaxation time of $\omega$

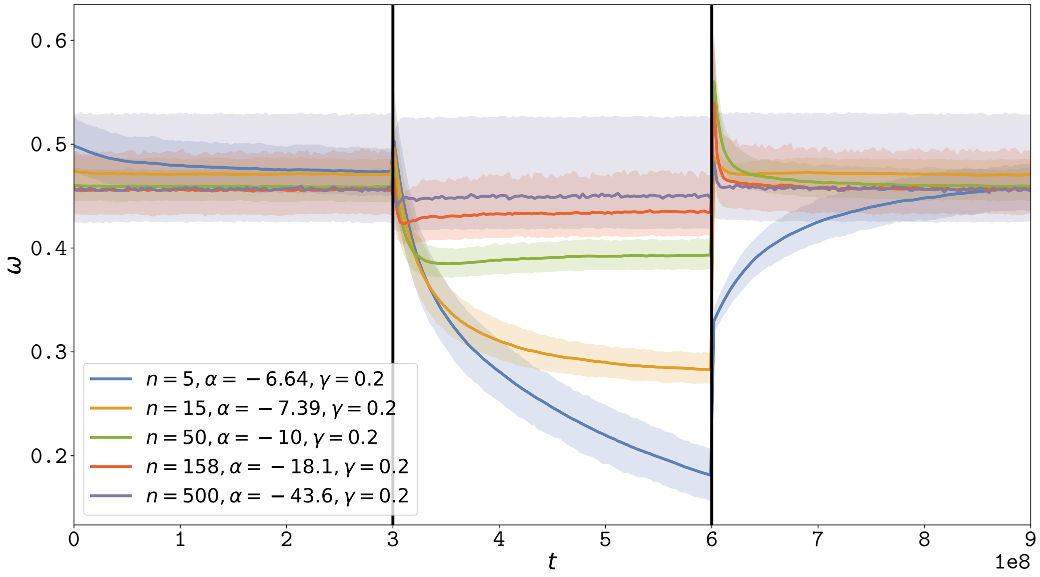

Figure 11: $\omega$ Relaxation after a brutal change in $N_\mathrm{e}$, for various $n$ while correcting for $\alpha$. The left and right panel correspond to low $N_\mathrm{e}$ ($1e^5$) and the middle panel corresponds to high $N_\mathrm{e}$ ($2e^6$). Solid line corresponds to the average over replicates ($r$) and the shaded area correspond to the 90% interval among replicates. The mutation rate ($\mu$) is $1e{-}8$ per year per site, and the total time of the computation is 900 million years. $\beta = 1.686$, $\gamma = 0.2$ for all simulations. The number of sites is changed from $n = 15$ to $n = 158$, and the number of replicates is changed accordingly such that the total number of sites ($n * r$) is kept constant. Moreover, $\alpha$ is changed according to $n$ and $\gamma$ such that the equilibrium value $x^*$ is kept constant, by solving numerically equation 18. Increasing $n$ implies a higher rate of relaxation.

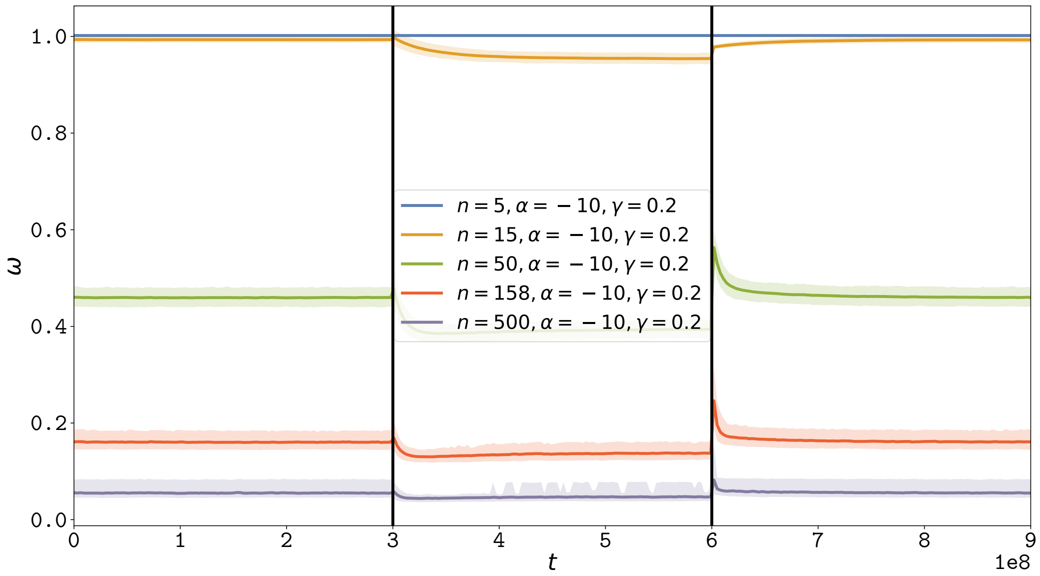

Figure 12: $\omega$ Relaxation after a brutal change in $N_e$, for various $n$. The left and right panel correspond to low $N_e$ ($1e^5$) and the middle panel corresponds to high $N_e$ ($2e^6$). Solid line corresponds to the average over replicates ($r$) and the shaded area correspond to the 90% interval among replicates. The mutation rate ($\mu$) is $1e{-8}$ per year per site, and the total time of the computation is 900 million years. $\beta = 1.686$, $\gamma = 0.2$ and $\alpha = -10$ for all simulations. The number of sites is changed from $n = 15$ to $n = 158$, and the number of replicates is changed accordingly such that the total number of sites ($n * r$) is kept constant. Increasing $n$ implies a higher $\omega$ at equilibrium, a lower response of the $\omega$ to changes in $N_e$ and a higher rate of relaxation.

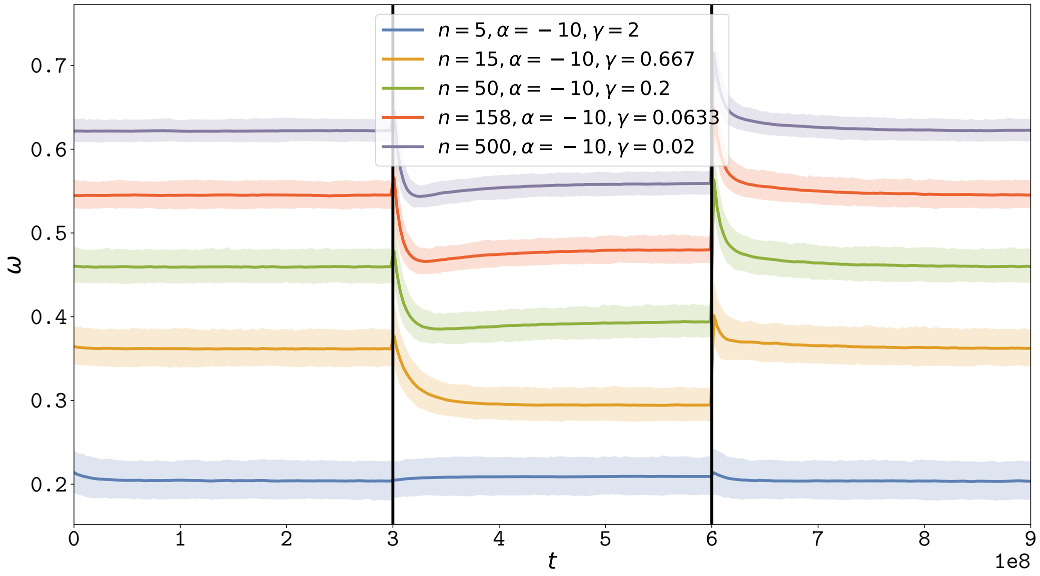

Figure 13: $\omega$ Relaxation after a brutal change in $N_e$, for various $n$ while correcting for $\gamma$. The left and right panel correspond to low $N_e$ ($1e^5$) and the middle panel corresponds to high $N_e$ ($2e^6$). Solid line corresponds to the average over replicates ($r$) and the shaded area correspond to the 90% interval among replicates. The mutation rate ($\mu$) is $1e{-8}$ per year per site, and the total time of the computation is 900 million years. $\beta = 1.686$, $\alpha = -10$ for all simulations. The number of sites is changed from $n = 15$ to $n = 158$, and the number of replicates is changed accordingly such that the total number of sites ($n * r$) is kept constant. Moreover, $\gamma$ is changed according to $n$ such that the product $\gamma n$ is kept constant, thus the response of the $\omega$ to changes in $N_e$ is kept constant. Increasing $n$ implies a higher $\omega$ at equilibrium, and a higher rate of relaxation.

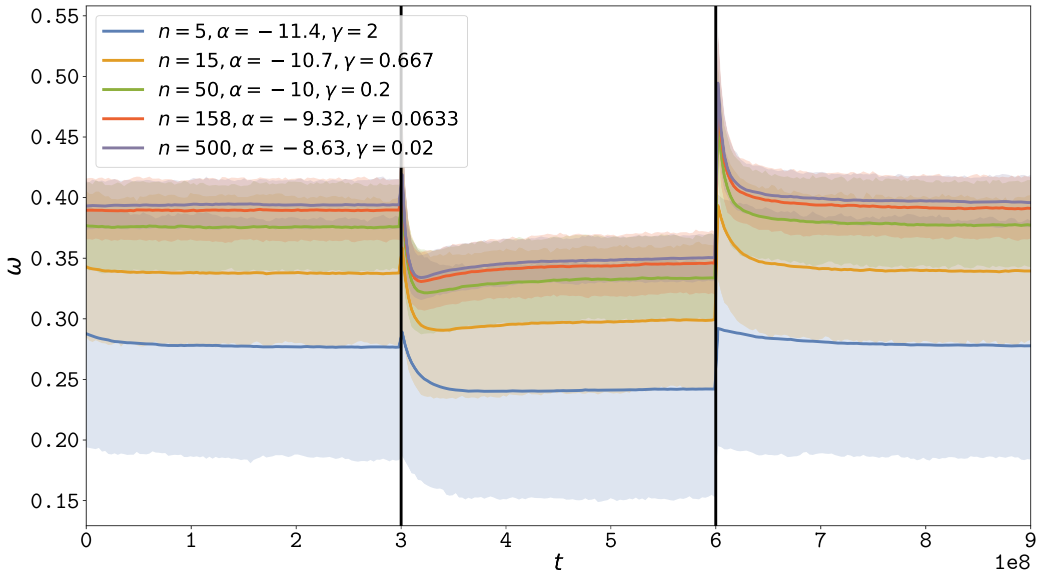

Figure 14: $\omega$ Relaxation after a brutal change in $N_{\mathrm{e}}$, under a Grantham model. The left and right panel correspond to low $N_{\mathrm{e}}$ ($1e^5$) and the middle panel corresponds to high $N_{\mathrm{e}}$ ($2e^6$). Solid line corresponds to the average over replicates ($r$) and the shaded area correspond to the 90% interval among replicates. The mutation rate ($\mu$) is $1e{-}8$ per year per site, and the total time of the computation is 900 million years. $\beta = 1.686$, $\gamma = -10$ for all simulations. The number of sites is changed from $n = 15$ to $n = 158$, and the number of replicates is changed accordingly such that the total number of sites ($n * r$) is kept constant. Moreover, $\gamma$ is changed according to $n$ such that the product $\gamma n$ is kept constant, thus the response of the $\omega$ to changes in $N_{\mathrm{e}}$ is kept constant. Finally, $\alpha$ is changed according to $n$ and $\gamma$ such that the equilibrium value $x^*$ is kept constant, by solving numerically equation 18. Increasing $n$ implies a higher rate of relaxation.

### 8 Distribution of fitness effects

DNA mutations changing a genotype can result in a change of phenotype, and ultimately a change in fitness. From a specific genotype, all the possible mutations thus result in a distribution of phenotypic effect (DPE) and fitness effects (DFE). The DPE and DFE are not known a priori, but are the resulting consequence of the mutation-selection-drift balance. Empirically, these distributions are of particular importance since they can be obtained experimentally or inferred with other data. As an example, DFE can be inferred from polymorphism dataset (Eyre-walker and Keightley, 2007; Galtier, 2016). Moreover, the distribution of $\Delta\Delta G$ for novel mutations can be obtained experimentally.

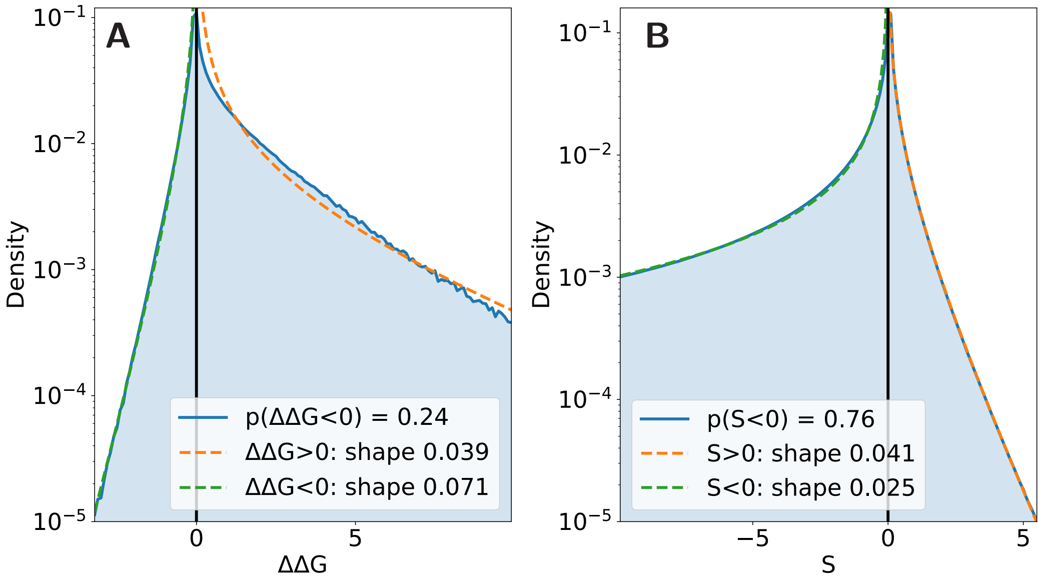

Figure 15: Distribution of fitness effects and phenotypic effect for novels non-synonymous mutations observed along a simulation at the mutation-selection balance. $\alpha = -118$, $\gamma = 1$, $n = 300$, $\beta = 1.686$, and for each non-optimal amino acid, $\gamma$ is scaled by the Grantham distance to the optimal amino acid. Each side of the distribution is fitted to a gamma distribution, shown in dotted line. Panel A. Distribution of observed $\Delta\Delta G$, which fit adequately the gamma distribution for negative $\Delta\Delta G$ (stabilizing mutations). Panel A. Distribution of observed selection coefficient, which fit adequately the gamma distribution for both positive and negative selection coefficient. However the shape parameter estimated is not the same for positive and negative selection coefficients.
